# The NEDD8 cycle controlled by NEDP1 upon DNA damage is a regulatory module of the HSP70 ATPase activity

**DOI:** 10.1101/583864

**Authors:** Aymeric P. Bailly, Aurelien Perrin, Marina Serrano-Macia, Chantal Maghames, Orsolya Leidecker, Helene Trauchessec, ML Martinez-Chantar, Anton Gartner, Dimitris P. Xirodimas

**Author notes:** Present address: Max Planck Institute for Biology of Ageing Joseph-Stelzmann-Str. 9b D-50931, Cologne, Germany (Orsolya Leidecker). Equal author contribution (Chantal Maghames, Orsolya Leidecker).

## Abstract

Ubiquitin and ubiquitin-like chains are finely balanced by the action of conjugating and de-conjugating enzymes. Alterations in this balance trigger signalling events required for the response to stress conditions and are often observed in pathologies. How such changes are detected is not well-understood. We show that upon DNA damage the induction of the de-NEDDylating enzyme NEDP1 restricts the formation of poly-NEDD8 chains, mainly through lysines K11/K48. This promotes APAF1 oligomerisation and apoptosis induction, a step that requires the HSP70 ATPase activity. We found that HSP70 binds to NEDD8 and *in vitro*, mono-NEDD8 stimulates the ATPase activity of HSP70, counteracted upon poly-NEDDylation. This effect is independent of NEDD8 conjugation onto substrates. The studies identify the HSP70 chaperone as sensor of changes in the NEDD8 cycle, providing mechanistic insights for a cytoplasmic role of NEDD8 in the DNA damage induced apoptosis. They also indicate that the balance between mono- versus poly-NEDDylation is a regulatory module of HSP70 function. The above findings may be important in tumorigenesis, as we find that NEDP1 levels are downregulated in Hepatocellular Carcinoma with concomitant accumulation of NEDD8 conjugates.

## Introduction

A key property of ubiquitin and ubiquitin-like (Ubl) molecules is their ability to modify substrates as single moieties or in the form of polymeric chains. The extent and topology of polymeric chains is finely balanced by the coordinated action of conjugating and de-conjugating enzymes (Williamson et al., 2013). The activity of these enzymes is altered as part of the cellular response to stress and is de-regulated in pathological conditions including cancer, immunological and neurodegenerative diseases (Popovic et al., 2014). Hence, these enzymes are regarded as major targets for therapeutic intervention. The outcome of such de-regulation is the change in the equilibrium between the mono- and polymeric state of ubiquitin and Ubl modification. However, the pathways that sense such alterations are not well-understood (Popovic et al., 2014; Williamson et al., 2013).

The ubiquitin-like molecule NEDD8 is highly conserved and essential in almost all tested organisms. Its functions have been characterised mainly through mono-NEDDylation of the cullin family of proteins and stimulation of the activity of Cullin-Ring-Ligases (CRL), but also through modification of non-cullin substrates (Abidi and Xirodimas, 2015; Enchev et al., 2014). Defects in the NEDD8 cycle resulting in increased levels of NEDDylation have been reported in several types of cancers including lung adenocarcinomas, squamous-cell carcinoma and hepatocellular carcinoma and inhibitors of the NEDD8 pathway are in Phase II clinical trials (Abidi and Xirodimas, 2015; Barbier-Torres et al., 2015; Delgado et al., 2018). Protein NEDDylation is a reversible process. The NEDP1 (DEN1, SENP8) protease specifically processes NEDD8 into the mature form, required for the activation of NEDD8 by the NEDD8 E1 enzyme and additionally catalyses de-NEDDylation of substrates (Abidi and Xirodimas, 2015; Enchev et al., 2014). While proteomic studies have indicated the formation of NEDD8 chains in cells, their regulation and biological function(s) are not well-defined (Abidi and Xirodimas, 2015; Enchev et al., 2014). By combining studies in *C. elegans* and in human cells we found a conserved role of the NEDD8 cycle in the DNA damage induced apoptosis. The induction of NEDP1 upon DNA damage restricts the formation of poly-NEDD8 chains mainly through lysines K11/K48 in the cytoplasm. This promotes the oligomerisation of the Apoptotic Protease Activating Factor 1 (APAF1) and apoptosis induction. We found that de-NEDDylation is required for the release of the HSP70 chaperone from APAF1, a required step towards APAF1 oligomerisation. HSP70 binds to NEDD8 and we mapped the ATPase domain as the binding site for NEDD8 on HSP70. Biochemical analysis shows that the balance between mono- and poly-NEDD8 is a regulatory module for HSP70 function; mono-NEDD8 activates the ATPase activity of HSP70, which is counteracted upon poly-NEDDylation. Restriction of poly-NEDDylation by NEDP1 restores the stimulatory effect of NEDD8 on HSP70 ATPase activity. The studies reveal that HSP70 is a sensor of changes in the NEDD8 cycle controlled by NEDP1 and provide mechanistic insights on the role of poly-NEDDylation restriction as regulatory module of HSP70 function and apoptosis induction upon DNA damage. These findings may be relevant in pathology, as we found that NEDP1 levels are downregulated in a mouse model system for Hepatocellular Carcinoma with concomitant accumulation of NEDD8 conjugates. Collectively, the data provide a molecular basis for a potential suppressive role of NEDP1 in tumorigenesis through restriction of poly-NEDD8 chains.

## Results

### The de-NEDDylating enzyme ULP-3/NEDP1 restricts the formation of K11/48 NEDD8 chains and is required for DNA damage-induced apoptosis in *C.elegans*

We identified *ulp-3* (<u>U</u>biquitin-like <u>p</u>rotease-<u>3</u>, sequence Y48A5A.2) as the *C. elegans* homologous gene of human NEDP1 by reciprocal BLAST analysis (Fig. 1A). ULP-3 has the catalytic triad His/Asp/Cys that defines the cysteine protease super-family (Fig. 1A). *In vitro*, ULP-3 processes the NEDD8 C-terminal similarly to NEDP1, whereas *in vivo*, overexpression of ULP-3 in human cancer cells decreases NEDDylation of L11, a previously characterised NEDD8 substrate (Xirodimas et al., 2008) and has no effect on total ubiquitination. Both activities depend of the predicted catalytic C167 (Supplemental information Fig. 1). These results indicate that *C. elegans* ULP-3 is a *bona fide* NEDD8 specific protease and the *C. elegans* homologous protein to human NEDP1.

**Figure 1.**
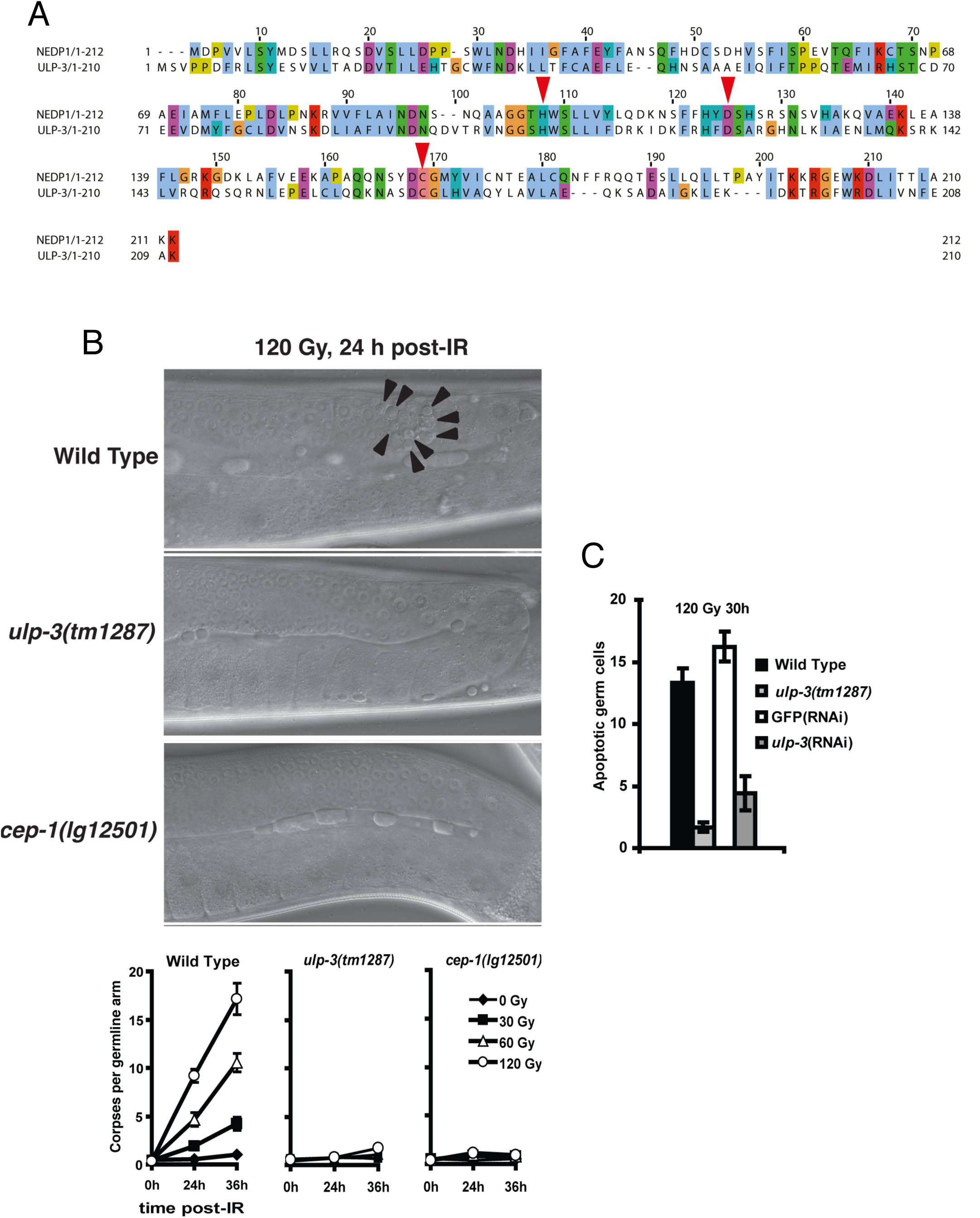
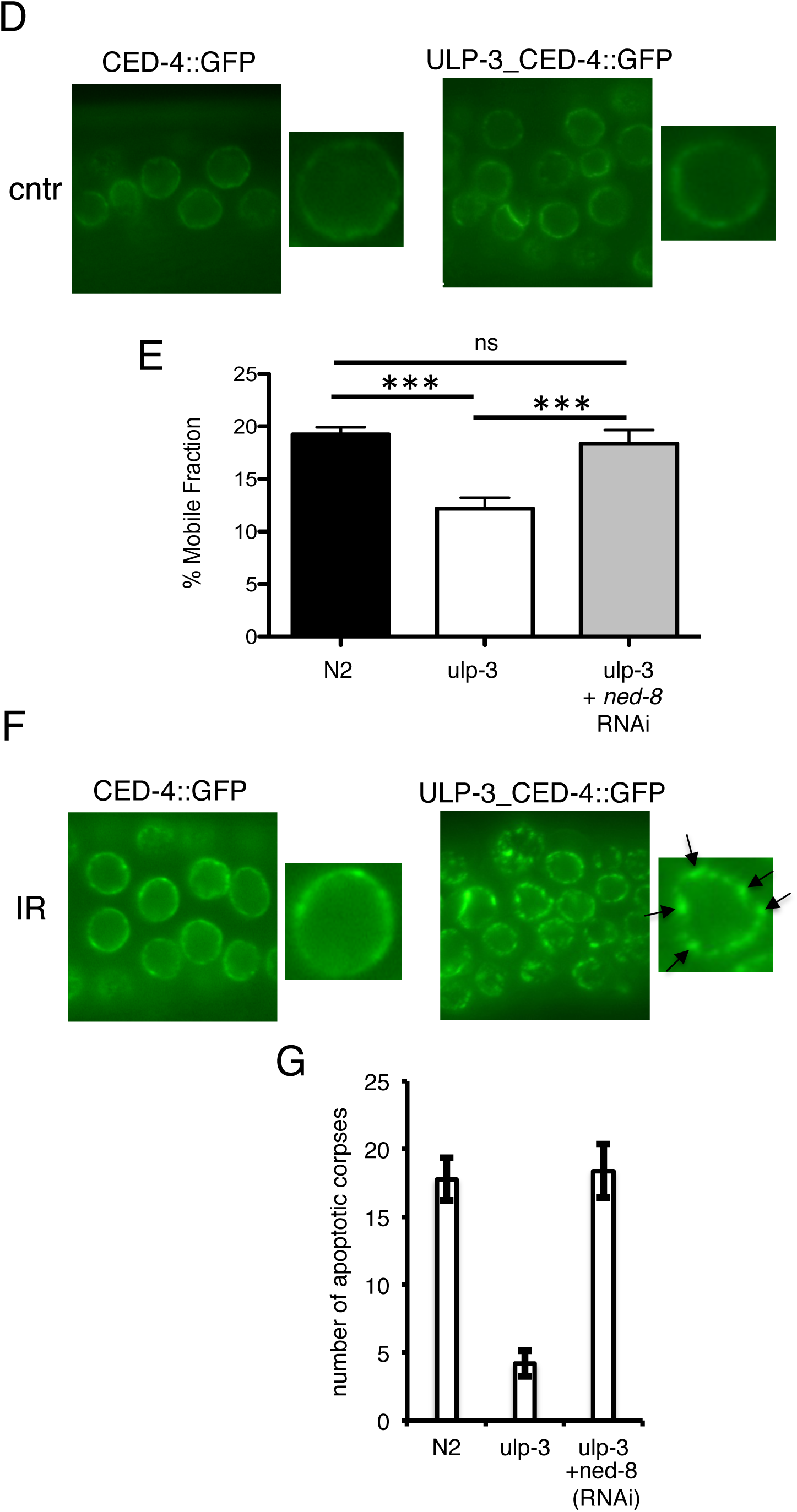
The de-NEDDylating enzyme ULP-3 is required for DNA damage-induced apoptosis in *C.elegans*. A. ClustalWS alignment of human NEDP1 and *C. elegans* ULP-3 shows the conservation of the catalytic triad His106/Asp123/Cys167 (red arrows). B. Wild type, *ulp-3* and *cep-1* deleted animals were irradiated as indicated. Germ cells were monitored through DIC optics and apoptotic cells were counted. The average number of apoptotic cells for at least 15 animals of each genotype is plotted with error bars including SEM (bottom panel). C. Similar experiment as in B, including RNAi treatments in wild type animals as indicated. D. CED-4::GFP localization in wild type and *ulp-3* mutant germ cells. Living worms were dissected and gonads immediately prepared for confocal microscopy. E. CED-4::GFP mobile fraction is determined by Fluorescence Recovery After Photobleaching (FRAP) in the indicated genetic backgrounds and RNAi treatment. Average values (*n*=20) of mobile fraction with error bars as SEM (t-test, p ≤ 0.001). F. CED-4::GFP localisation in the indicated backgrounds 24hrs after 120Gy IR. Arrows indicate the CED-4::GFP punctate structures at the perinuclear area in *ulp-3* mutant germ cells. G. *ned-8* RNAi in *ulp-3* deleted animals. Average number of apoptotic cells was determined as in B (*n*>15) with SEM as error bars.

We obtained a deletion allele (*ulp-3 (tm1287))*, within the reading frame, which abolishes *ulp-3* expression, shown by RNA and protein level analysis (Supplemental information Fig. 2). The *ulp-3* KO animals are viable and further systematic phenotypic characterisation shows no defects in cell cycle progression, growth and fertility compared to wild type animals (data not shown). However, worms deleted or silenced for *ulp-3*, in contrast to wild type animals, are almost completely resistant to the induction of apoptosis upon ionising radiation (IR) in germ cells (Fig. 1B, C). Therefore, ULP-3 is not essential for viability and development in *C. elegans* but it is required for the IR-induced apoptosis.

**Figure 2.**
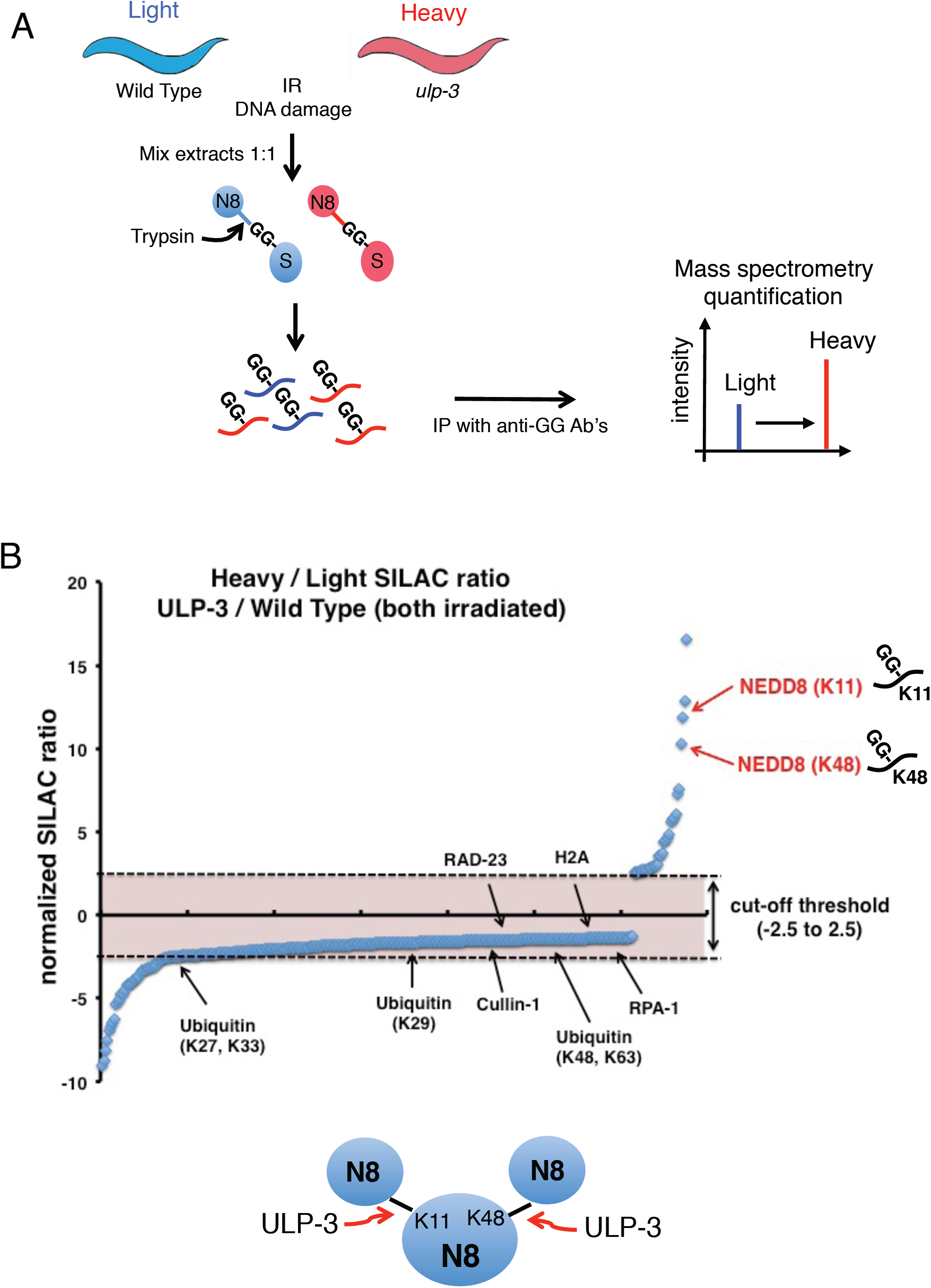
ULP-3 restricts the formation of K11/K48 NEDD8 chains. A. Proteomic SILAC strategy in *C.elegans* to identify ULP-3 targets upon DNA damage. Worms labelled with light (wild type) or heavy (*ulp-3* mutant) isotopic amino acids were exposed to 120Gy of IR. Protein extracts were mixed 1:1 ratio and after trypsin digestion diGly containing peptides were immunoprecipitated and analysed by MS. B. Plot of normalised SILAC ratio of unique diGly peptides (top panel). Diagram for the role of ULP-3 in restricting NEDD8 chain formation through K11/K48 (right panel).

The apoptotic core pathway in *C. elegans* is responsible for both germ cells homeostasis and developmental programmed cell death (Bailly and Gartner, 2013; Lettre and Hengartner, 2006). We exploited the worm mutant *ced-1 (e1735)* in which apoptotic corpses engulfment is severely impaired, allowing quantitation of developmental apoptosis. No difference in the number of apoptotic cells that persist in the head of L1 larvae between the *ced-1* single mutant with the *ulp-3; ced-1* double mutant was observed, suggesting a specific role for ULP-3 in germ cells apoptosis upon IR (Supplemental information Fig. 3). By using the ts *glp-4* mutant in which the germ line is eliminated at the restrictive temperature or by analysing dissected germ lines, we found by qPCR and western blot analysis that *ulp-3* is preferentially expressed in germ cells, providing an explanation for the specific role of ULP-3 in the IR-induced apoptosis in germ cells (Supplemental information Fig. 4).

**Figure 3.**
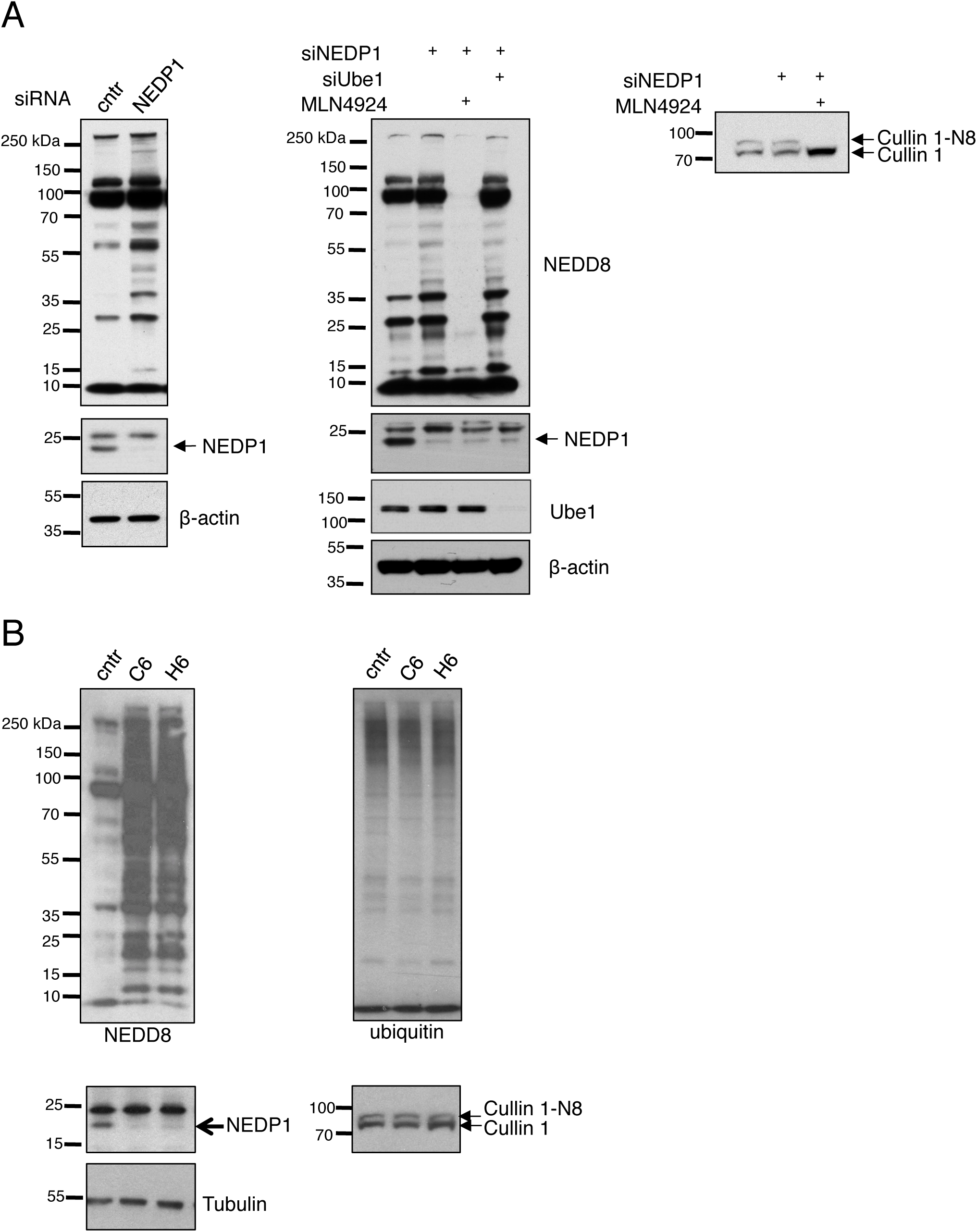
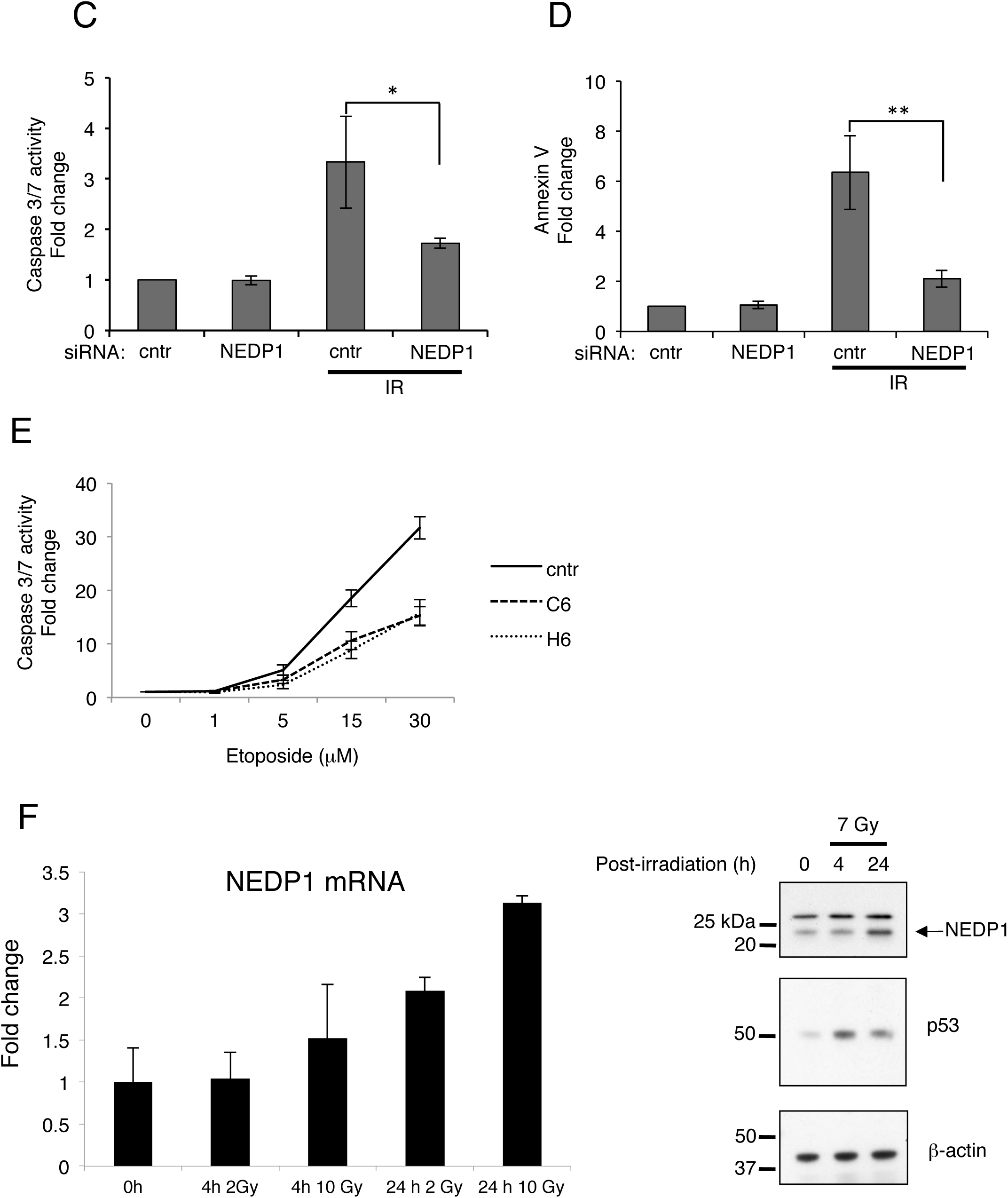
Conserved role for NEDP1 in the DNA damage-induced apoptosis. A. Western blotting of extracts from U2OS cells transfected with siRNAs and treated with the NEDD8 E1 inhibitor MLN4924 (1μM, 15hrs) as indicated. Extracts were re-analysed with cullin-1 antibodies (right panel). B. Western blot analysis of extracts from parental U2OS or U2OS NEDP1 knockout clones (C6, H6). C, D. Similar experiment as in A, except cells were treated with IR (7Gy) and analysed either for caspase 3/7 activity (C) or AnnexinV staining (D) 9hrs later. E. Caspase 3/7 activity in parental or NEDP1 knockout U2OS cells exposed to etoposide for 15hrs. Average values (*n*=3) with SEM as error bars. F. MCF7 cells were irradiated with 2 or 10Gy for the indicated times. Quantitative real-time PCR for NEDP1 was carried out as described in Methods. The experiments were performed in triplicates; Average values +/-SEM (left panel). Western blot analysis in extracts of MCF7 cells treated as indicated (right panel).

**Figure 4.**
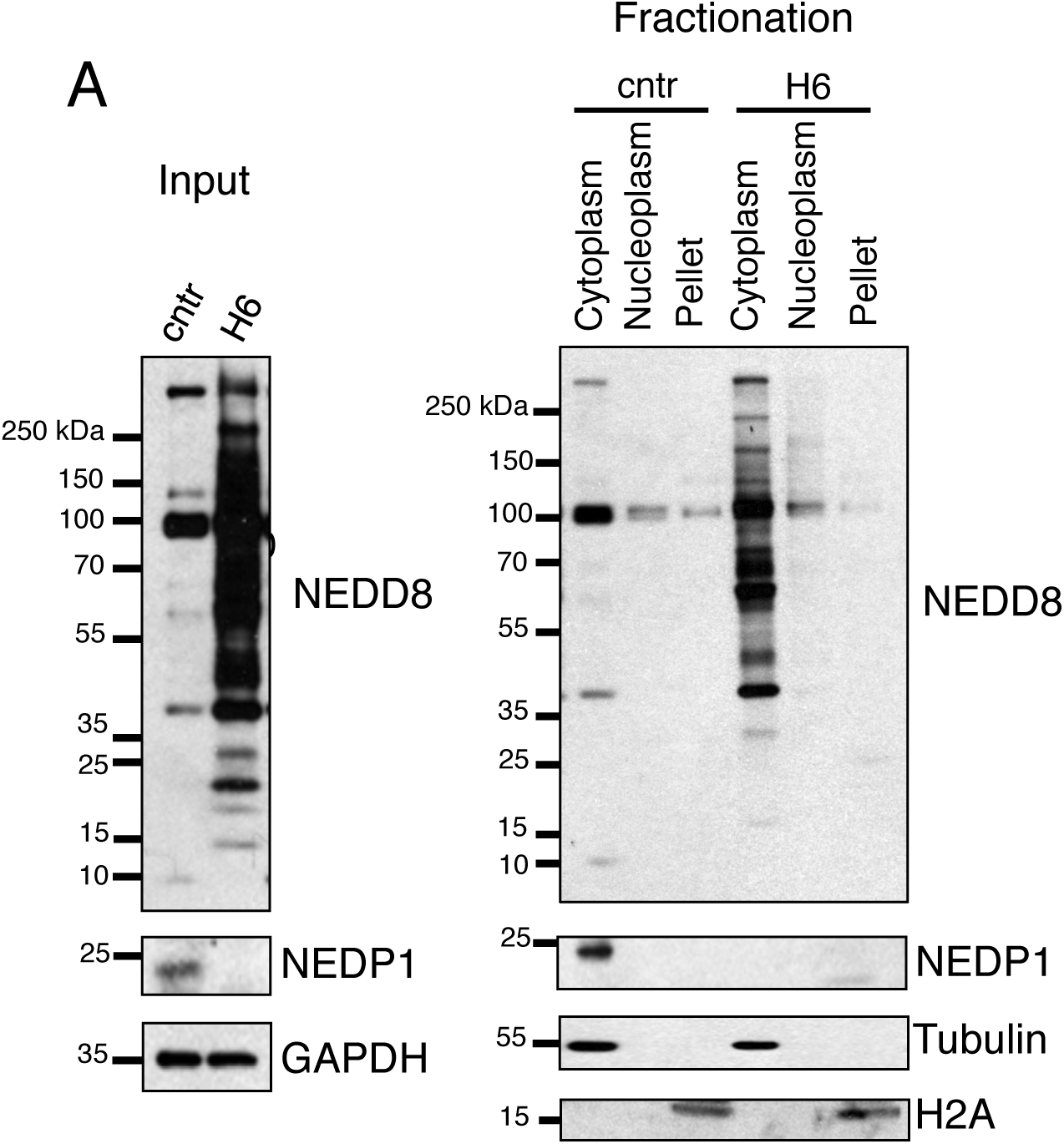
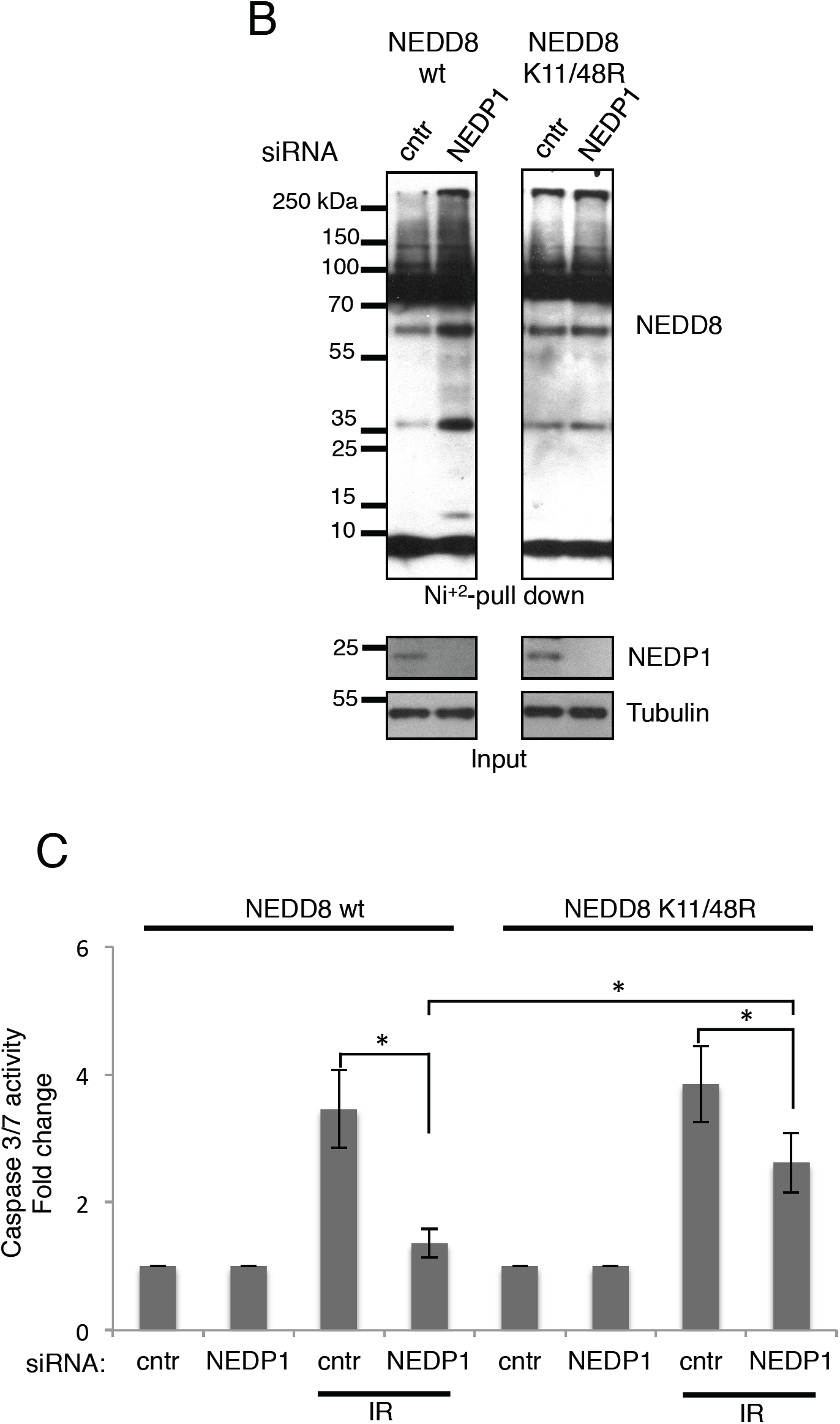
Restriction of K11/K48 NEDD8 chains in the cytoplasm is required for DNA-damage induced apoptosis. A. Subcellular fractionation in parental or H6 NEDP1 knockout U2OS cells and western blotting for the indicated proteins. B. U2OS cells stably expressing either wild type or K11/48R His_6_-NEDD8 mutant were transfected with the indicated siRNAs. Isolated His_6_-NEDD8 conjugates were blotted for NEDD8. Expression levels of indicated proteins in total cell extracts (Input) C. Similar experiment as in Fig. 3C. Average values (*n*=3) with SEM as error bars. p values *≤ 0.05, **≤ 0.01.

In response to Double Strand Breaks (DSBs), the main cellular circuitry involved in apoptosis induction is the DNA damage checkpoint signalling pathway that ultimately triggers the highly conserved transcription activator p53, called CEP-1 in *C. elegans (<u>C</u>. <u>e</u>legans* <u>p</u>53-like <u>1</u>)(Bailly and Gartner, 2013; Lettre and Hengartner, 2006). We analysed multiple phenotypes indicative of an efficient activation of the DNA damage checkpoint signalling pathway: We determined the sensitivity of the *ulp-3* mutant to IR in a clonogenic survival assay indicative of defects in DSBs repair, assessed the cell cycle arrest induced by DNA damage and quantitatively measured the induction of the two CEP-1 target genes, *egl-1* and *ced-13*, as readout of CEP-1/p53 activation. *ulp-3* deletion has no effect on DNA repair, cell cycle arrest and CEP-1 activation upon IR (Supplemental information Fig. 5) suggesting that the cascade of events that follow the activation of the DNA damage checkpoint signalling pathway are not under ULP-3 regulation. We concluded that the branching point where ULP-3 intersects the DNA damage checkpoint signalling pathway, is downstream or parallel to the *C. elegans* p53-like protein CEP-1.

**Figure 5.**
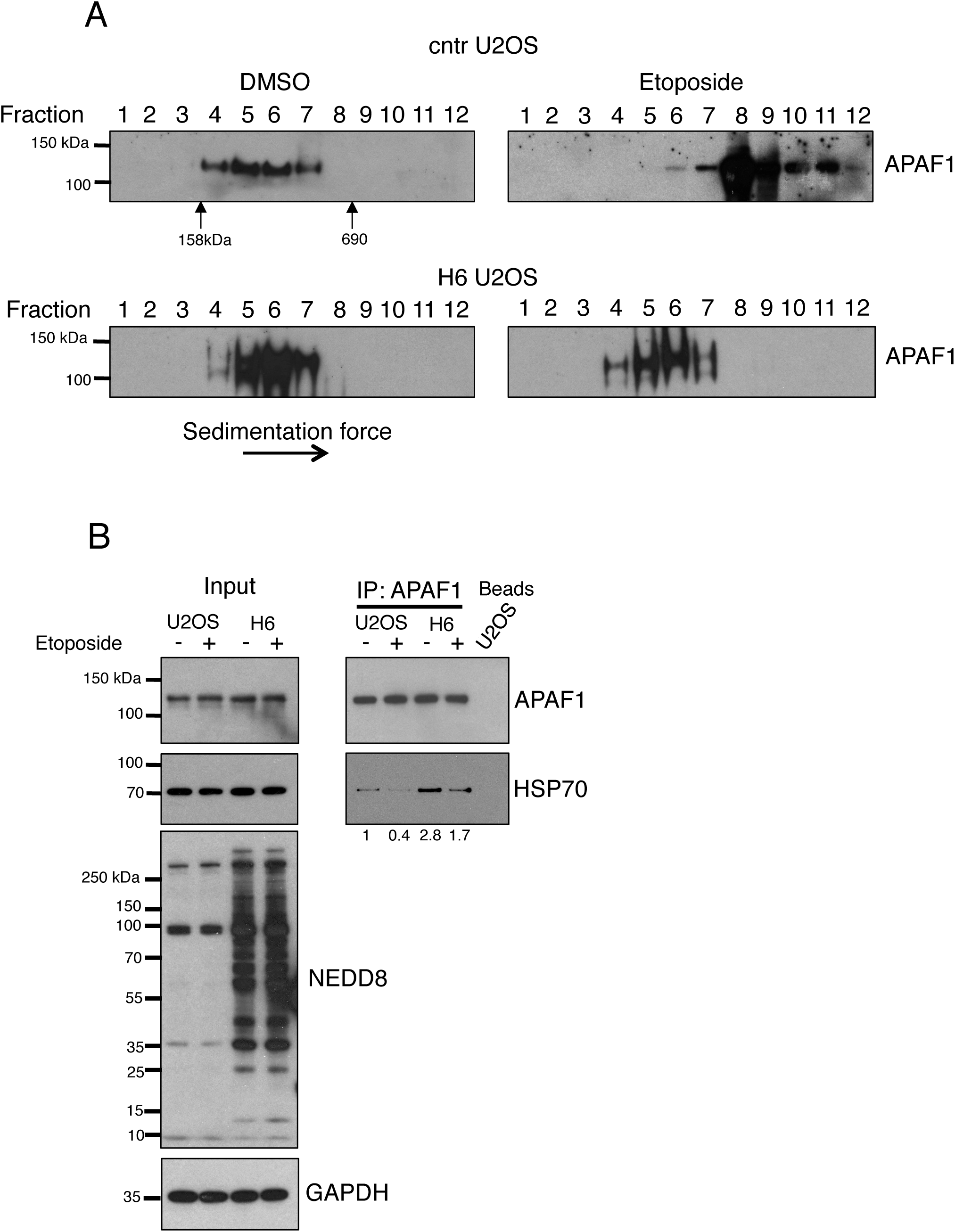
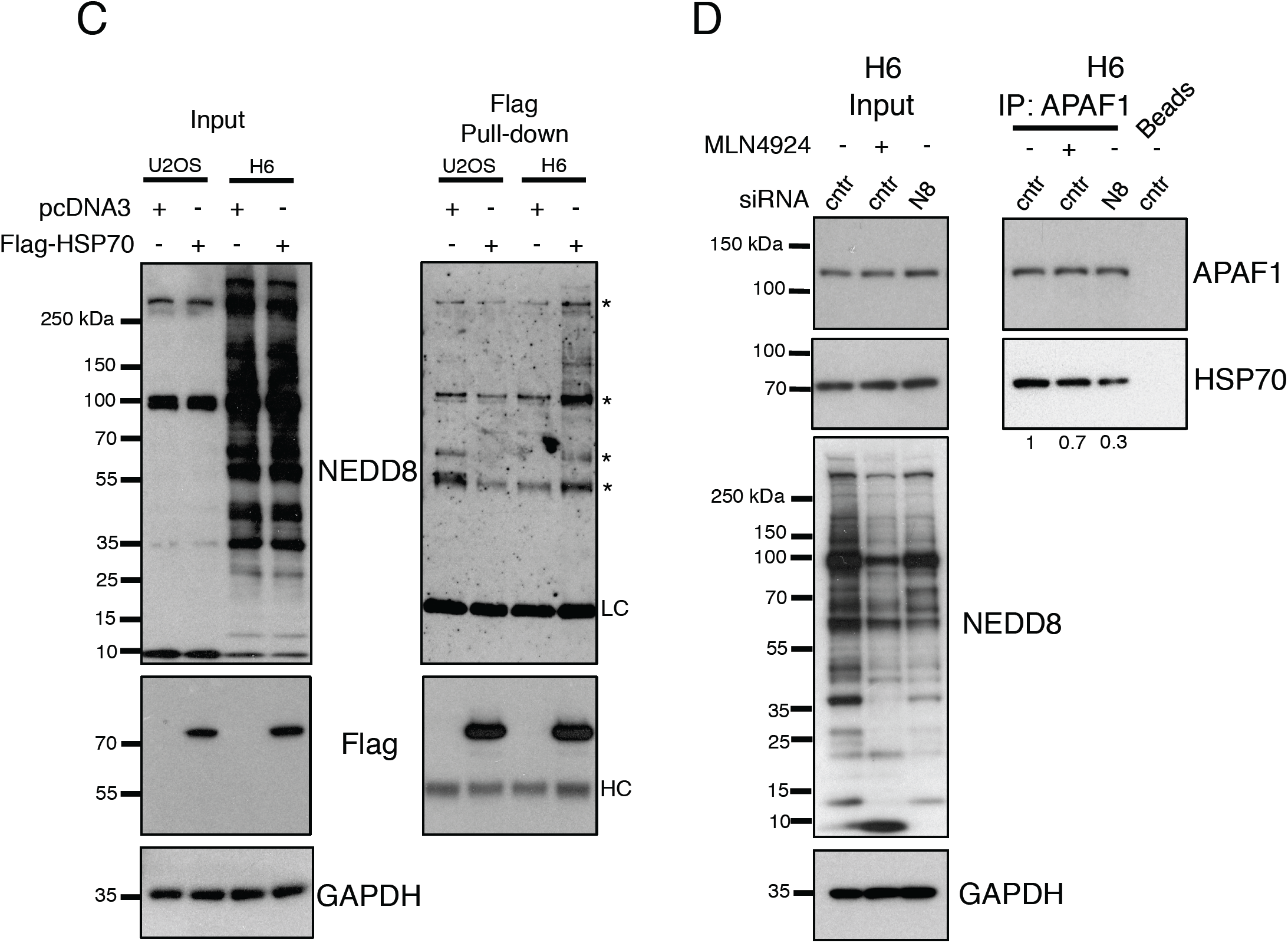
NEDP1 is required for HSP70-mediated APAF1 oligomerisation upon DNA damage. A. Parental or H6 cells were either untreated (DMSO) or exposed to etoposide (50μM) for 15hrs before extracts were subjected to sucrose gradient centrifugation. Fractions were analysed by western blotting for APAF1. Arrows indicate the separation of Aldolase (158kDa) and thyroglobulin (690kDa). B. Parental or H6 cells were treated as in (A) and extracts were used for immunoprecipitation (IP) with APAF1 antibodies. Total cell extracts (Input) and IPs were analysed by western blotting with the indicated antibodies. C. Parental or H6 NEDP1 KO U2OS cells stably expressing 3xFlag-HSP70 were used for immunoprecipitations and western blotting as indicated. Asterisks indicate non-specific bands. LC, HC: Light and Heavy chain respectively. D. H6 cells were transfected with control (cntr) or NEDD8 (N8) siRNAs for 48hrs. MLN4924 (1μM) was added for 15hrs. Signals for HSP70 were quantified with Image Gauge.

Downstream of CEP-1 activation, the oligomerisation of CED-4 (APAF1 homologue) into the apoptosome homomeric complex is the critical step for caspase 3-like protein (CED-3) activation and apoptosis induction (Qi et al., 2010). We monitored the effect of *ulp-3* deletion on CED-4 localisation using a *C. elegans* strain stably expressing CED-4::GFP under its endogenous promoter and 3’UTR. CED-4::GFP displays perinuclear localisation consistent with the pattern of endogenous CED-4 (Fig. 1D). By using <u>F</u>luorescence <u>R</u>ecovery <u>A</u>fter <u>P</u>hotobleaching (FRAP) in living worms we determined the mobile fraction of CED-4::GFP in the perinuclear area, as 19.24 % ± 0.69 %, which is compatible with the mobile fraction of a membranous or membrane interacting protein (Wolter et al., 1997). Deletion of *ulp-3* significantly reduces the CED-4::GFP mobile fraction by nearly 40% (12.18% ± 1.03%) (Fig. 1D, E). DNA damage by IR lead to the appearance of CED-4::GFP as punctate structures at the vicinity of the nuclear membrane, and this phenomenon was exacerbated in the *ulp-3* mutant (Fig. 1F). FRAP analysis showed that these CED-4::GFP structures at the perinuclear membrane become completely immobile (data not shown). The observed defects in the CED-4 mobile fraction and apoptosis induction upon IR in the *ulp-3* mutant were restored to a nearly wild type level upon transient *ned-8* RNAi treatment (Fig. 1E, G). The data indicate that de-NEDDylation by ULP-3 increases the mobile fraction of CED-4 required for the activation of the apoptotic response to IR.

To gain mechanistic insights into the role of ULP-3 in the IR-induced apoptosis, we devised an unbiased proteomics approach to discover potential NEDD8 targets for ULP-3 upon DNA damage. Identification of diglycine (diGly) remnants left on lysine residues upon trypsin digestion of proteins by mass spectrometry demonstrates their modification with ubiquitin, NEDD8 or ISG15 (Kessler, 2013; Ordureau et al., 2015). We hypothesised that changes in the diGly signature upon deletion of the de-NEDDylating enzyme ULP-3, should specifically indicate changes in the NEDD8 modification repertoire. We combined Stable Isotope Labelling with Amino acid in nematodes (SILAC) (Larance et al., 2011) with the use of antibodies that recognise the diGly remnant on modified peptides (Kessler, 2013; Ordureau et al., 2015; Xu et al., 2010). Wild type and *ulp-3* deleted worms were labelled with Light and Heavy isotopes respectively (Fig. 2A). To specifically address the effect of ULP-3 on the NEDD8 proteome upon DNA damage, both sets of worms were exposed to IR. Extracts were mixed in 1:1 ratio before immunoprecipitation with the anti-diGly antibody followed by MS/MS (Fig. 2A). A total of 951 non-redundant diGly peptides were quantified, among which the abundance of 32 was significantly increased (>2.5 fold) upon *ulp-3* deletion after IR (Supplemental information Table 1). Interestingly, no significant changes in the modification of multiple proteins involved in the DNA damage response, including cullins, were observed (Fig. 2B). Similarly, ubiquitin chain formation was mainly unaffected under these conditions (Fig. 2B). The proteomic data are consistent with the genetic characterisation, suggesting that ULP-3 does not control the DNA damage checkpoint activation. By contrast, we observed a significant increase for the modification of NEDD8 on lysines 11 and 48 (K11/K48, SILAC ratio of 10.6 and 11.2 respectively) (Fig. 2B), indicating that deletion of ULP-3 increases poly-NEDD8 chain formation. The data highlight that a key function of ULP-3 upon DNA damage is to restrict the formation of NEDD8 chains through lysines K11/K48 (Fig. 2B).

### NEDP1 is a DNA damage induced gene with a conserved role in DNA damage induced apoptosis in human cells

To determine and biochemically explore the conservation of our findings in *C. elegans*, we investigated in human cells the role of the ULP-3 homologue NEDP1 in the DNA damage-induced apoptosis. Knockdown of NEDP1 by siRNA increased overall protein NEDDylation with no significant effect on cullin-NEDDylation (Fig. 3A). Importantly, the observed increase in NEDDylation depends on the canonical NEDD8 activating enzyme (NAE) and not on NEDD8 activation by the ubiquitin E1 enzyme Ube1, which occurs under proteotoxic stress (Hjerpe et al., 2012; Leidecker et al., 2012) (Fig. 3A, middle panel). A dramatic increase in protein NEDDylation with no effect on ubiquitination is observed in 2 different clones of U2OS (C6 and H6) where NEDP1 is deleted using CRISPR/Cas9 (Fig. 3B, Supplemental information Fig. 6). By monitoring caspase 3/7 activation or Annexin V staining, we found that NEDP1 knockdown significantly decreases the IR-induced apoptosis (Fig. 3C, D). Similar results were obtained using the NEDP1 KO U2OS cells exposed to the chemical inducer of DSBs, etoposide (Fig. 3E). Under these stress conditions, NEDP1 deletion does not affect either γ H2AX or p53 stabilisation (Supplemental information Fig. 7). Importantly, DSBs induce a late (24hrs post IR) expression of the NEDP1 mRNA and protein levels, showing that NEDP1 is a DNA damage responsive gene (Fig. 3F). Consistent with the notion that NEDP1 does not control early events in the DNA damage response at the chromatin level, subcellular fractionation shows that both NEDP1 and the accumulated NEDD8 conjugates upon NEDP1 deletion are almost exclusively localised in the cytoplasm (Fig. 4A). These findings are consistent with the observations in *C. elegans* showing that the upstream activation of the DNA damage checkpoint is unaffected by ULP-3 deletion. Based on the proteomic analysis in *C. elegans* we tested in human cells the role of K11/48 NEDD8 chain formation in the induction of apoptosis upon DSBs. We generated U2OS cells stably expressing either wild type or K11/48R His_6_-NEDD8 mutant at endogenous levels (Liu and Xirodimas, 2010) (Fig. 4B). As expected in the presence of wild type NEDD8, knockdown of NEDP1 increases NEDDylation and prevents the induction of apoptosis upon DSBs (Fig. 4B, C). However, the K11/48R NEDD8 mutant impaired the increase in NEDDylation upon NEDP1 knockdown and partially restored (up to 70%) the IR-induced apoptosis (Fig. 4B, C). In combination with the proteomic analysis in *C. elegans*, the data suggest that the de-NEDDylation of poly-NEDD8 chains via their K11/48 linkage is the main and conserved function of ULP-3/NEDP1 that is required for the induction of apoptosis upon DNA damage.

**Figure 6.**
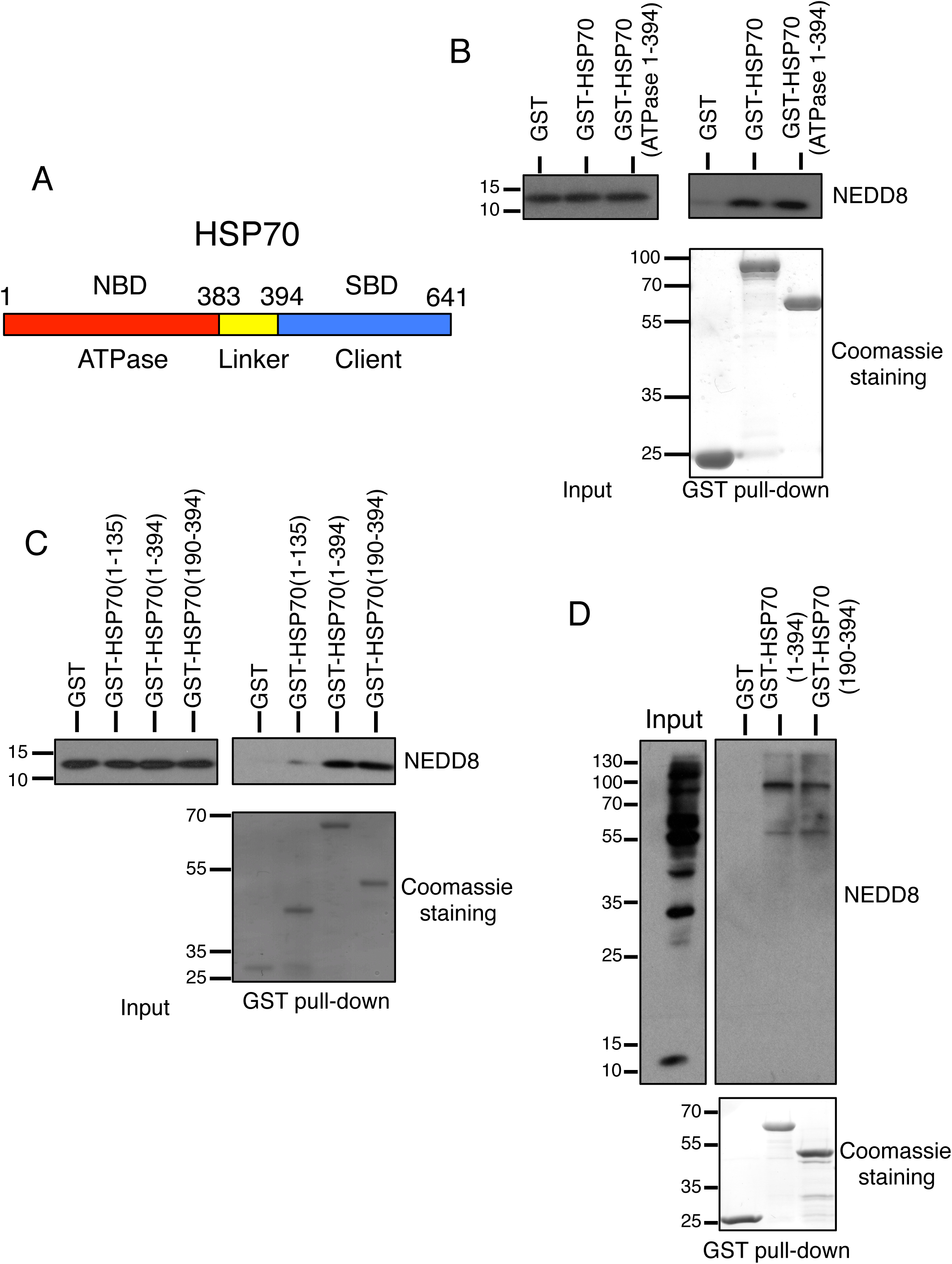
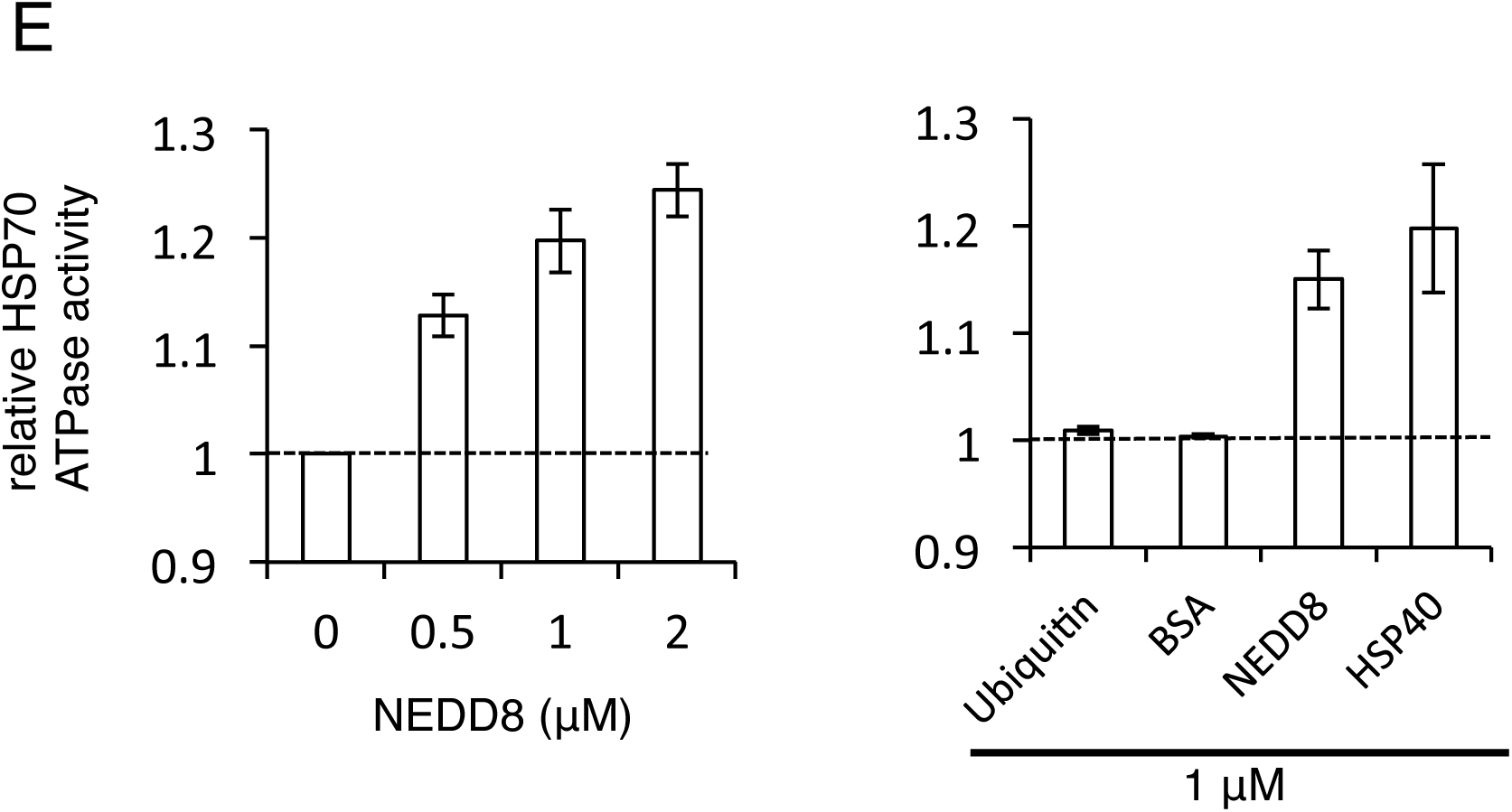
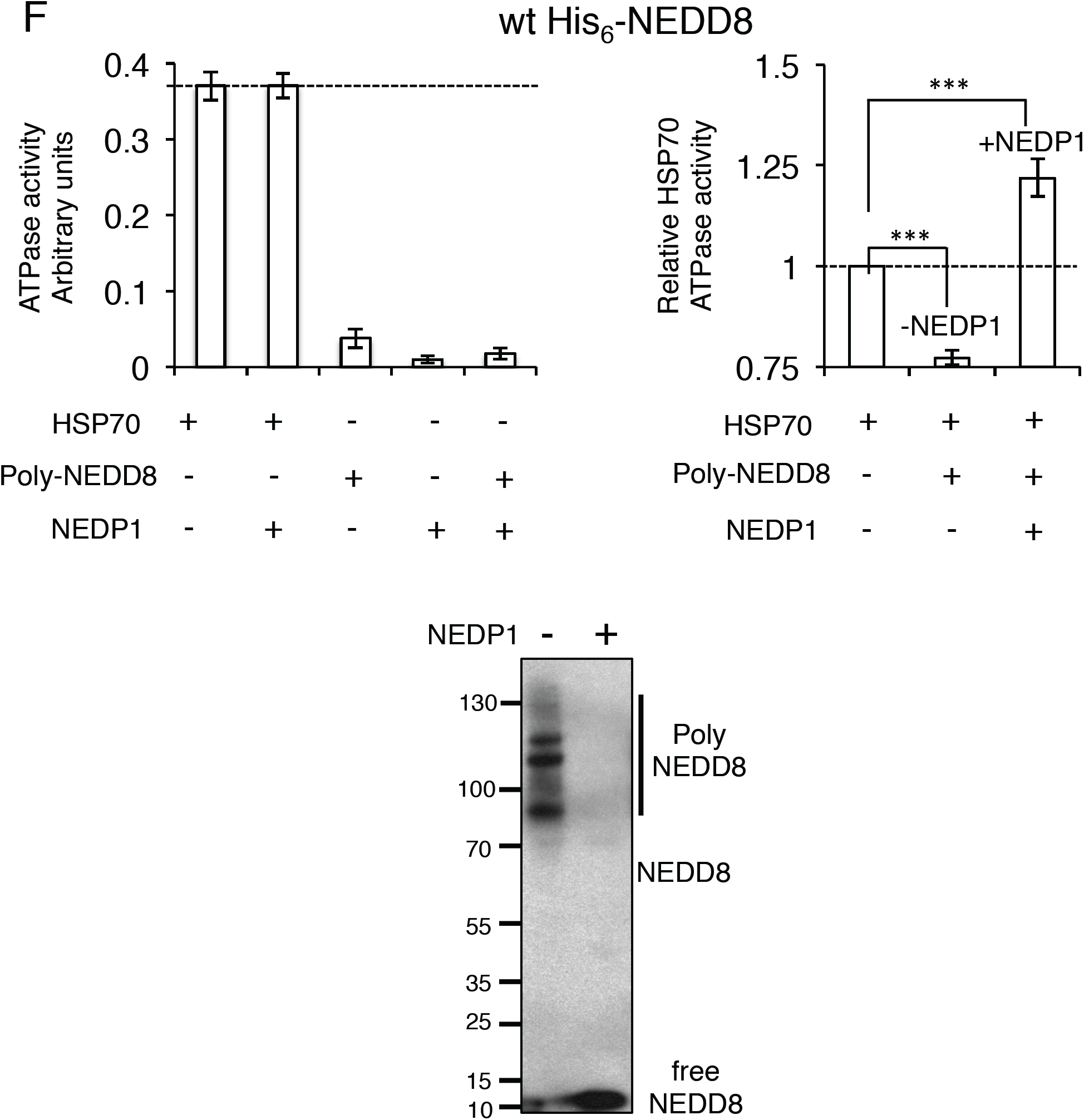
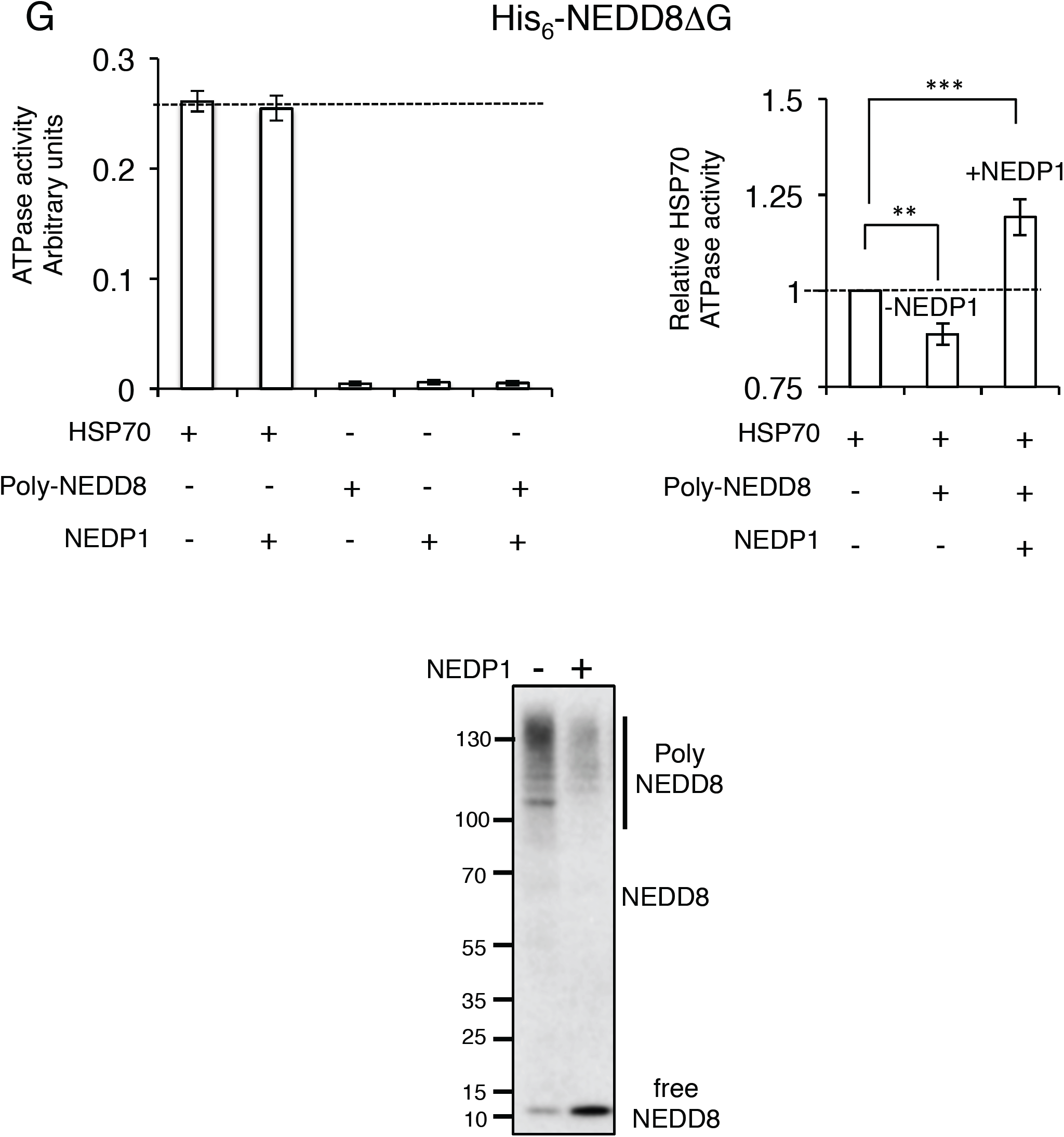
Mono-NEDD8 stimulates the ATP hydrolysis activity of HSP70, which is compromised upon poly-NEDDylation. A. Representation of the Nucleotide (NBD) and Substrate (SBD) Binding Domains in HSP70. B, C. Recombinant GST or various indicated GST-HSP70 constructs were incubated with recombinant NEDD8. Input and GST pull-downs were analysed by western blotting with anti-NEDD8 antibody. D. Cells extracts from H6 NEDP1 KO cells were incubated with GST, GST-HSP70 (1-394) or GST-HSP70 (190-394). Input and GST pull-downs were analysed by western blotting with anti-NEDD8 antibody. E. The stimulation of the ATP hydrolysis activity of HSP70 was measured in the presence of increasing concentrations of NEDD8 (*n*=3 +/-SEM) (left). The relative stimulation of HSP70 ATPase activity by NEDD8 was compared to that of HSP40 (DNAJB4); Ubiquitin and BSA were used as negative controls (*n*=3 +/- SEM) (right). F. Poly-NEDD8 conjugates were isolated from H6 NEDP1 KO cells stably expressing His_6_-NEDD8 by Ni^2+^ pull-down. Poly-NEDD8 conjugates were subjected to NEDP1 digestion (bottom panel) and tested in a HSP70 ATP hydrolysis activity assay (right panel, *n*=5 +/-SEM). Neither poly-NEDD8 conjugates nor recombinant NEDP1 carry contaminant ATP hydrolysis activity (left graph). G. Similar experiment as in (F) with the exception that H6 NEDP1 knockout U2OS cells were transfected with the His_6_-NEDD8ΔG (*n*=5 +/- SEM), p values **≤ 0.01, ***≤ 0.001.

**Figure 7.**
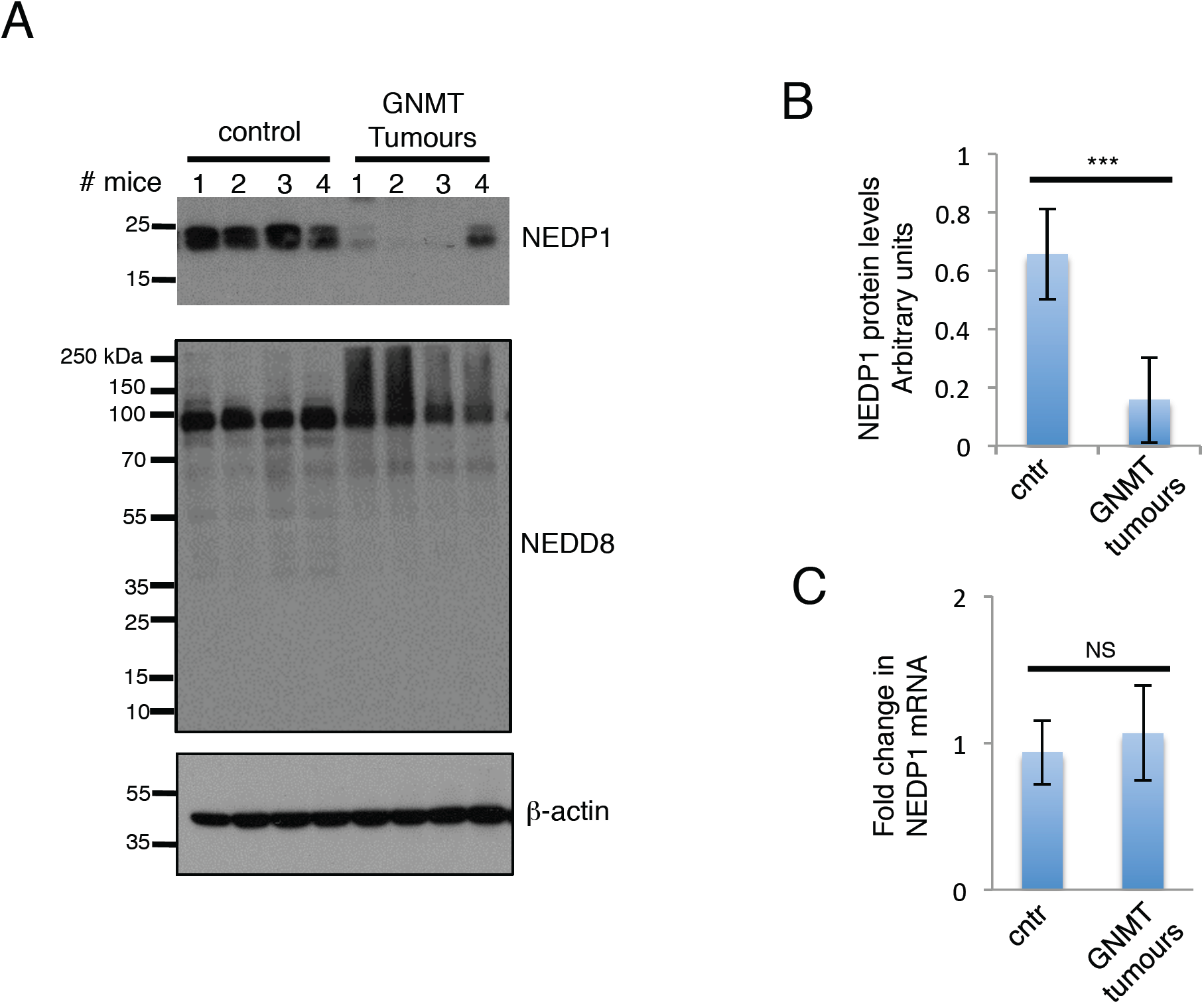
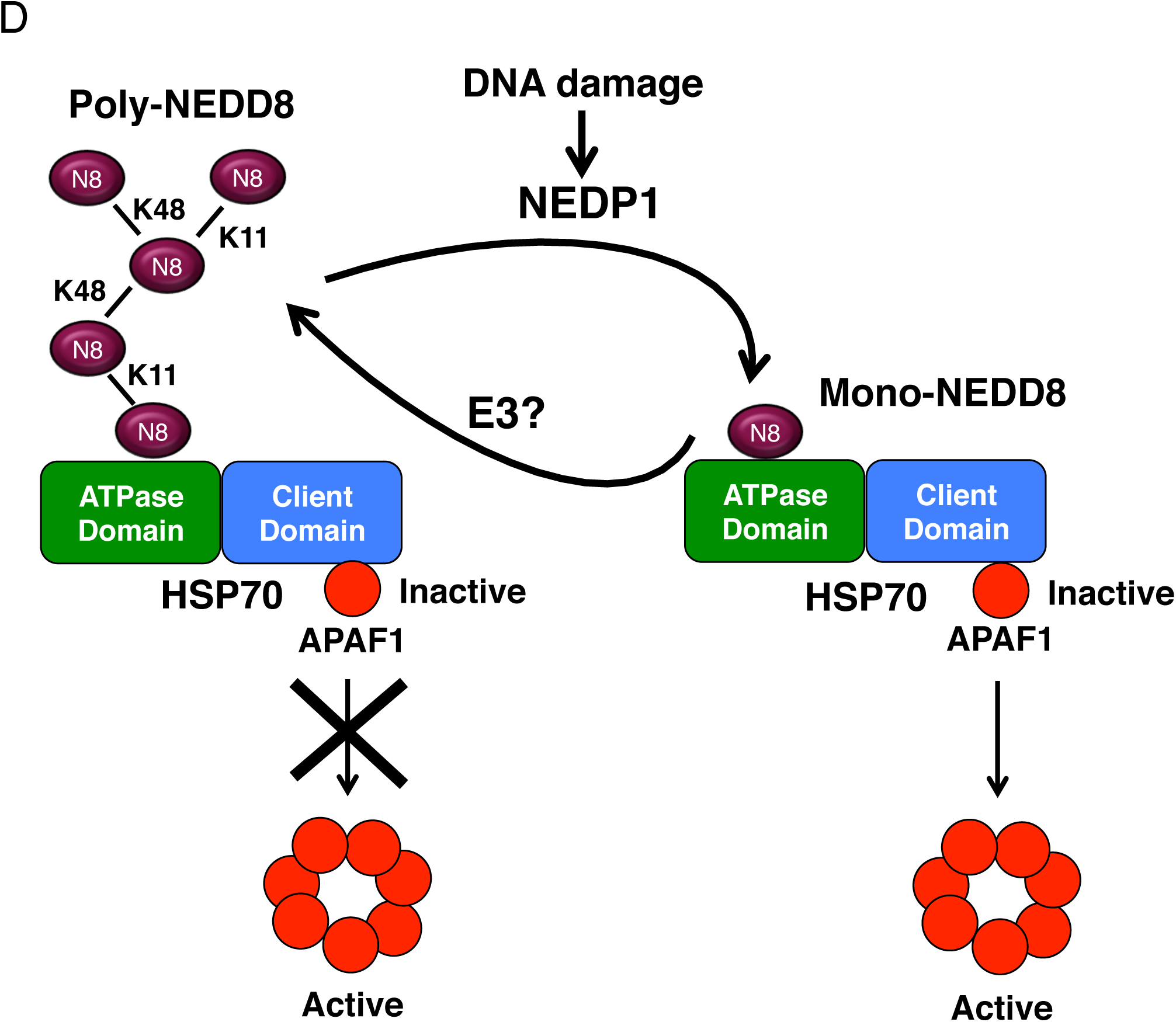
NEDP1 levels are downregulated in Hepatocellular Carcinoma. A. Liver tissue extracts from control or GNMT knockout tumoral group mice were analysed for the levels of NEDP1 and NEDD8 by western blotting. B-actin was used as loading control. B. Quantitation of NEDP1 protein levels shown in (A). Data show the average of the NEDP1/β-actin ratio of 4 tested samples +/- SEM. p values ***≤ 0.001. C. Quantitative real-time PCR for nedp1 mRNA was carried out as described in Methods. Values represent the average of 8 independent experiments for each condition +/- SEM. D. Model for the role of the NEDD8 cycle in HSP70 activity regulation and apoptosis induction upon DNA damage. Poly-NEDD8 chains interact with HSP70 and prevent the release and oligomerisation of APAF1 into an active complex. The induction of NEDP1 upon DNA damage restricts poly-NEDD8 chains into mono-NEDD8, which activates HSP70 ATPase activity resulting in APAF1 oligomerisation and activation. The NEDD8 cycle controlled by NEDP1 acts as a regulatory module for HSP70 function.

### NEDP1 is required for HSP70-mediated APAF1 oligomerisation upon DNA damage

Based on the findings in *C. elegans* on the role of ULP-3 on CED-4 mobility, we followed a biochemical approach in human cells to determine the role of NEDP1 on APAF1 activation upon DSBs. We monitored by sucrose gradient sedimentation the oligomerisation of APAF1, indicative of the formation of the apoptosome (Bratton and Salvesen, 2010). While in control cells DSBs cause the formation of APAF1 oligomers, in NEDP1 KO cells APAF1 oligomerisation was severely impaired and APAF1 remained in its monomeric state (Fig. 5A). The findings in *C. elegans* and human cells indicate that ULP-3/NEDP1 through de-NEDDylation promotes the formation of the active apoptosome upon DSBs.

APAF1 oligomerisation is a dynamic and key event towards initiation of apoptosis (Bratton and Salvesen, 2010; Cain et al., 1999; Rodriguez and Lazebnik, 1999). Amongst several control mechanisms, the binding of the heat shock protein 70 (HSP70) to APAF1 is an important regulatory event in APAF1 oligomerisation. More specifically, the activation and release of HSP70 from APAF1 upon stress, is required for the apoptosome formation and induction of apoptosis (Beere et al., 2000; Saleh et al., 2000). However, the signals, mechanism(s) and co-chaperone(s) involved in HSP70 activation under these conditions remain unclear.

Consistent with the previous studies (Beere et al., 2000; Saleh et al., 2000) we found that DSBs caused the decrease of HSP70 binding to APAF1 in control U2OS cells (Fig. 5B). However, in the NEDP1 knockout H6 cells an increased HSP70-APAF1 binding was observed both in unstressed and DSBs stress conditions (Fig. 5B). We investigated several possibilities of how the increase in NEDDylation observed in NEDP1 KO cells impacts on the HSP70-APAF1 interaction, including assays monitoring the potential NEDDylation of HSP70 and APAF1. Under endogenous levels of NEDD8 expression and denaturing conditions that prevent non-covalent interactions, we did not detect the covalent modification of either HSP70 or APAF1 with NEDD8. In contrast, we found as the most consistent observation the interaction of HSP70 with the accumulated NEDD8 conjugates in H6 NEDP1 KO cells (Fig. 5C). This suggests that the binding of HSP70 with poly-NEDDylated conjugates in NEDP1 KO cells blocks the HSP70 release from APAF1. Consistent with this hypothesis, inhibition of poly-NEDDylation in NEDP1 KO cells, either with the specific NEDD8 E1 inhibitor MLN4924 or with NEDD8 siRNA, reduced the binding of HSP70 to APAF1 (Fig. 5D).

### NEDD8 binds to the ATPase domain of HSP70 and stimulates HSP70 ATPase activity

We then followed a biochemical approach to further characterise the NEDD8-HSP70 binding. Using recombinant proteins we found a direct interaction between NEDD8 and HSP70 *in vitro* (Fig. 6A, B). Mutational analysis showed that NEDD8 binds the ATPase domain of HSP70, between amino acids 190-394, which overlaps with the HSP40 J-domain binding region within the HSP70 ATPase domain (Fig. 6B, C) (Ahmad et al., 2011). Furthermore, the HSP70 (190-394) fragment can also pull-down NEDD8 conjugates from NEDP1 KO cells (Fig. 6D). While the *in vitro* analysis shows that HSP70 interacts with mono-NEDD8, the *in vivo* analysis suggests a preference of HSP70 for high molecular weight poly-NEDD8 conjugates.

ATP hydrolysis by HSP70 is a critical regulatory element for the interaction of HSP70 with client proteins which is stimulated by HSP70 co-chaperones, including HSP40 (Kityk et al., 2012; Mayer and Bukau, 2005; Palleros et al., 1991; Zhuravleva et al., 2012). We found that mono-NEDD8 but not ubiquitin stimulates the ATPase activity of HSP70 to a similar extent as HSP40, indicating an important role for NEDD8 for HSP70 function regulation (Fig. 6E, Supplemental information Fig. 8). Mutation of a single amino acid (I44A) within the hydrophobic patch of NEDD8 prevented both the NEDD8-HSP70 interaction and the stimulatory effect of NEDD8 on HSP70 ATPase activity (Supplemental information Fig. 9A, B).

### Poly-NEDDylation blocks the stimulatory effect of mono-NEDD8 on HSP70 ATPase activity

We next determined the role of poly-NEDDylation in HSP70 ATPase function. We isolated poly-NEDD8 conjugates either from NEDP1 knockout cells stably expressing His_6_-NEDD8 (Fig. 6F) or from *in vitro* reactions using recombinant His_6_-NEDD8 (Supplemental information Fig. 9C). We employed high-salt purification conditions to ensure that no contaminant ATPase activity was co-purified (Fig. 6F). In contrast to mono-NEDD8, the addition of poly-NEDD8 conjugates did not stimulate the HSP70 ATPase activity; rather a reproducible decrease was observed (Fig. 6F, Supplemental information Fig. 9C). However, upon de-conjugation with the addition of recombinant NEDP1, an increase in HSP70 ATPase activity was observed (Fig. 6F, Supplemental information Fig. 9C). We performed a similar experiment as in Fig. 6F, transfecting instead NEDP1 knockout cells with a His_6_-NEDD8 mutant deleted of its C-terminal glycine. This mutant is deficient in modifying substrates but itself can be modified by endogenous wild type NEDD8, allowing the isolation of NEDD8 chains that are not attached to substrates (unanchored). Similarly to the data obtained with the wild type NEDD8 chains, unanchored NEDD8 chains inhibited HSP70 ATPase activity, which was induced upon de-conjugation by NEDP1 (Fig. 6G). This shows the direct role of NEDD8 chain processing by NEDP1 on HSP70 ATPase activity regulation, which is independent on the conjugation of NEDD8 to substrates.

### NEDP1 is downregulated in Hepatocellular Carcinoma

The NEDD8 pathway is upregulated in several types of tumors, which often acquire resistance to the used chemotherapy (Abidi and Xirodimas, 2015). Based on our observations the accumulation of poly-NEDD8 chains upon NEDP1 deletion, compromises the induction of apoptosis upon DNA damage. However, the patho-physiological relevance for these findings remains unknown. We determined the expression of NEDP1 in the Glycine N-methyltransferase (GNMT) knockout pre-clinical mouse model of Hepatocellular Carcinoma (HCC). GNMT is a tumor suppressor and deletion in mice results in the development of HCC (Martínez-Chantar et al., 2008). We chose this system as previous studies showed upregulation in protein NEDDylation in *GNMT-/-* derived HCC tumors (Delgado et al., 2018). We found that NEDP1 protein levels in liver tissue extracts are dramatically reduced in all tested HCC animals compared to control animals (Fig. 7A, B). The observed decrease is not due to changes in *nedp1* mRNA levels suggesting a post-transcriptional mechanism (Fig. 7C). Importantly, the decrease in NEDP1 levels is accompanied with a significant increase in NEDDylation, particularly for the high molecular weight conjugates, indicative of poly-NEDD8 conjugates (Fig. 7A). The data indicate that decrease in NEDP1 levels is induced during liver tumorigenesis and provide a molecular mechanism for the reported increase of NEDDylation in these tumors.

## Discussion

The role of ubiquitin and SUMO modification in the DNA damage response is established at the chromatin level, where upon recruitment they initiate complex signalling events for DNA repair and/or the induction of apoptosis (Jackson and Durocher, 2013; Zhao et al., 2014). Nuclear functions of NEDD8 required for DNA repair have also been reported through NEDDylation of cullins and non-cullin targets, including histones H2A, H4 and PCNA (Li et al., 2014; Ma et al., 2013; Guan et al., 2017). In this study, the combination of *C. elegans* genetics and biochemistry in human cells reveals a previously uncharacterized cytoplasmic role for NEDD8, as protein quality control pathway of the apoptosome formation during the DNA-damage induced apoptosis.

In addition to the de-conjugating activity, NEDP1 also acts as a NEDD8 specific C-terminal hydrolase, exposing the di-glycine motif before NEDD8 activation by NAE (Abidi and Xirodimas, 2015; Enchev et al., 2014). The NEDD8 pathway is essential both in *C. elegans* and in humans so the absence of lethality upon NEDP1 deletion indicates the presence of additional hydrolases that can process NEDD8. While NEDP1 is reported to control the mono-NEDDylation of both cullin and non-cullin substrates (Aoki et al., 2013; Broemer et al., 2010; Coleman et al., 2017; Mergner et al., 2015; Watson et al., 2010; Xirodimas et al., 2008), the presented analysis shows that the restriction of poly-NEDD8 chains in the cytoplasm mainly through lysines K11/K48, is a key activity for ULP-3/NEDP1 that is required for the induction of apoptosis upon DNA damage.

Initiation of the apoptotic process relies on the assembly of the apoptosome, which comprises several proteins including APAF1. Oligomerisation of APAF1 provides the scaffold for the association and activation of caspases for apoptosis execution (Beere and Green, 2001). Our studies suggest that NEDP1 dependent de-NEDDylation promotes, at least partially, the release of HSP70 from APAF1, which is regarded as an important step towards APAF1 oligomerisation (Beere et al., 2000; Saleh et al., 2000). Association of HSP70 with its binding partners is mainly controlled by the N-terminal ATPase activity of HSP70. The intrinsic ATPase activity of HSP70 is low but it is stimulated by interacting co-chaperones, which increase the turnover of HSP70 binding to the substrate (Mayer and Bukau, 2005; Young, 2010). The biochemical analysis indicates that mono-NEDD8 has characteristics of an HSP70 co-chaperone as: 1) NEDD8 interacts within the same region in the ATPase domain of HSP70 as the J-domain of the HSP40 co-chaperone (Fig. 6C); 2) NEDD8 stimulates the ATPase activity of HSP70 to a similar extent as HSP40 (Fig. 6E); 3) The stimulatory effect of NEDD8 depends on the presence of the so-called linker domain in HSP70, a typical characteristic of co-chaperones (Jiang et al., 2007) (Supplemental information Fig. 8). However, upon NEDD8 polymerisation the stimulatory effect of NEDD8 on HSP70 ATPase activity is severely compromised, despite the interaction of HSP70 with poly-NEDD8 conjugates. Thus, the balance between mono- and poly-NEDD8 chains is a regulatory element for HSP70 function. Future structural studies should provide detailed insights for the role of NEDD8 in HSP70 ATPase activity stimulation and how this effect is compromised upon NEDD8 polymerisation.

The biochemical analysis proposes a mechanism for the role of the NEDD8 cycle on APAF1 oligomerisation and DNA damage induced apoptosis; the induction of NEDP1 upon DNA damage restricts poly-NEDD8 conjugates into mono-NEDD8, creating the stimulatory signal for the HSP70 ATPase activity and apoptosome formation (Fig. 7D). Consistent with this notion, accumulation of poly-NEDD8 conjugates upon deletion of NEDP1 compromises the release of HSP70 from APAF1 and subsequently APAF1 oligomerisation, direct biological outcomes of defective HSP70 function. An interesting arising model is the role of HSP70 as sensor of the NEDD8 cycle. The extent and possibly the topology of poly-NEDDylation controlled by NEDP1 may indeed act as a “rheostat” to finely tune HSP70 chaperone activity (Fig. 7D). The studies support a paradigm where NEDP1 acts as the molecular link to functionally connect the NEDD8 cycle with the HSP70 chaperone machinery upon DNA damage.

While we can not exclude that the NEDDylation of a yet unidentified target(s) is part of the presented signaling cascade, the *in vitro* data show that de-polymerisation of NEDD8 chains is sufficient to activate HSP70 independently of a NEDD8 substrate (unanchored NEDD8 chains). Unanchored ubiquitin chains have been reported to control several processes, including innate immune response, viral uncoating and aggresome formation (Banerjee et al., 2014; Hao et al., 2013; Ouyang et al., 2012; Zeng et al., 2010). The presented biochemical data on the role of unanchored NEDD8 chains on HSP70 function along with recent studies proposing substrate independent functions for NEDD8 in the control of poly(ADP-ribose) polymerase 1 (PARP-1) activity (Keuss et al., 2019), suggest that unanchored NEDD8 chains can act as regulatory signals.

The observed downregulation of NEDP1 levels in HCC indicates that NEDP1 is a target during tumorigenesis. This finding defines a molecular basis for the reported defects in the NEDD8 cycle in tumors as evidenced by an increase in protein NEDDylation (Abidi and Xirodimas, 2015). Importantly, our findings on the role of NEDP1 in the DNA damage induced apoptosis through regulation of the HSP70-APAF1 module, provide potential mechanistic insights for the reported resistance of advanced HCC tumors to chemotherapy (Abidi and Xirodimas, 2015; Delgado et al., 2018). As inhibitors for the NEDD8 pathway are in Phase II clinical trials (Abidi and Xirodimas, 2015), NEDP1 levels may provide a marker for the tumor response to these inhibitors.

## Acknowledgements

We are grateful to the Montpellier RIO Imaging facility. The project is funded by the “ATIP/AVENIR” fellowship, the Labex EpiGenMed, an « Investissements d’avenir » program, reference ANR-10-LABEX-12-01, INCa (PLBIO16-251), the Marie Curie FP7 Re-integration programme and the Ministerio de Economía, Industria y Competitividad: SAF2017-87301-R (M.L.M-C.), BIOEF (Basque Foundation for Innovation and Health Research: EiTB Maratoia BIO15/CA/016/BD (M.L.M.-C). AB is recipient of the “ANR retour” postdoctoral fellowship and MS is recipient of the Asociación Española contra el Cáncer predoctoral fellowship.

## Author contribution

AB and AP performed all experiments in *C. elegans*. AB performed the biochemical analysis on HSP70-NEDD8 interaction. HT, CM and OL performed the experiments in human cells. MSM designed and performed all experiments in mice. DPX, AG and AB conceived and designed the experiments. DPX with the help of all authors wrote the manuscript.

## References

Abidi, N., and Xirodimas, D.P. (2015). Regulation of cancer-related pathways by protein NEDDylation and strategies for the use of NEDD8 inhibitors in the clinic. Endocr. Relat. Cancer 22, T55–T70.

Ahmad, A., Bhattacharya, A., McDonald, R.A., Cordes, M., Ellington, B., Bertelsen, E.B., and Zuiderweg, E.R.P. (2011). Heat shock protein 70 kDa chaperone/DnaJ cochaperone complex employs an unusual dynamic interface. Proc. Natl. Acad. Sci. U. S. A. 108, 18966–18971.

Aoki, I., Higuchi, M., and Gotoh, Y. (2013). NEDDylation controls the target specificity of E2F1 and apoptosis induction. Oncogene 32, 3954–3964.

Bailly, A., and Gartner, A. (2013). Germ cell apoptosis and DNA damage responses. Adv. Exp. Med. Biol. 757, 249–276.

Bailly, A.P., Freeman, A., Hall, J., Déclais, A.-C., Alpi, A., Lilley, D.M.J., Ahmed, S., and Gartner, A. (2010). The Caenorhabditis elegans Homolog of Gen1/Yen1 Resolvases Links DNA Damage Signaling to DNA Double-Strand Break Repair. PLoS Genet. 6, e1001025.

Banerjee, I., Miyake, Y., Nobs, S.P., Schneider, C., Horvath, P., Kopf, M., Matthias, P., Helenius, A., and Yamauchi, Y. (2014). Influenza A virus uses the aggresome processing machinery for host cell entry. Science 346, 473–477.

Barbier-Torres, L., Delgado, T.C., García-Rodríguez, J.L., Zubiete-Franco, I., Fernández-Ramos, D., Buqué, X., Cano, A., Gutiérrez-de Juan, V., Fernández-Domínguez, I., Lopitz-Otsoa, F., et al. (2015). Stabilization of LKB1 and Akt by neddylation regulates energy metabolism in liver cancer. Oncotarget 6, 2509–2523.

Beere, H.M., and Green, D.R. (2001). Stress management - heat shock protein-70 and the regulation of apoptosis. Trends Cell Biol. 11, 6–10.

Beere, H.M., Wolf, B.B., Cain, K., Mosser, D.D., Mahboubi, A., Kuwana, T., Tailor, P., Morimoto, R.I., Cohen, G.M., and Green, D.R. (2000). Heat-shock protein 70 inhibits apoptosis by preventing recruitment of procaspase-9 to the Apaf-1 apoptosome. Nat. Cell Biol. 2, 469–475.

Bratton, S.B., and Salvesen, G.S. (2010). Regulation of the Apaf-1-caspase-9 apoptosome. J. Cell Sci. 123, 3209–3214.

Brenner, S. (1974). The genetics of Caenorhabditis elegans. Genetics 77, 71–94.

Broemer, M., Tenev, T., Rigbolt, K.T.G., Hempel, S., Blagoev, B., Silke, J., Ditzel, M., and Meier, P. (2010). Systematic in vivo RNAi analysis identifies IAPs as NEDD8-E3 ligases. Mol. Cell 40, 810–822.

Cain, K., Brown, D.G., Langlais, C., and Cohen, G.M. (1999). Caspase activation involves the formation of the aposome, a large (approximately 700 kDa) caspase-activating complex. J. Biol. Chem. 274, 22686–22692.

Coleman, K.E., Békés, M., Chapman, J.R., Crist, S.B., Jones, M.J., Ueberheide, B.M., and Huang, T.T. (2017). SENP8 limits aberrant neddylation of NEDD8 pathway components to promote cullin-RING ubiquitin ligase function. eLife 6.

Delgado, T.C., Barbier-Torres, L., Zubiete-Franco, I., Lopitz-Otsoa, F., Varela-Rey, M., Fernández-Ramos, D., and Martínez-Chantar, M.-L. (2018). Neddylation, a novel paradigm in liver cancer. Transl. Gastroenterol. Hepatol. 3, 37–37.

Enchev, R.I., Schulman, B.A., and Peter, M. (2014). Protein neddylation: beyond cullin–RING ligases. Nat. Rev. Mol. Cell Biol. 16, 30–44.

Guan, J., Yu, S., and Zheng, X. (2017). NEDDylation antagonizes ubiquitination of proliferating cell nuclear antigen and regulates the recruitment of polymerase η in response to oxidative DNA damage. Protein Cell.

Hao, R., Nanduri, P., Rao, Y., Panichelli, R.S., Ito, A., Yoshida, M., and Yao, T.-P. (2013). Proteasomes Activate Aggresome Disassembly and Clearance by Producing Unanchored Ubiquitin Chains. Mol. Cell 51, 819–828.

Hjerpe, R., Thomas, Y., Chen, J., Zemla, A., Curran, S., Shpiro, N., Dick, L.R., and Kurz, T. (2012). Changes in the ratio of free NEDD8 to ubiquitin triggers NEDDylation by ubiquitin enzymes. Biochem. J. 441, 927–936.

Jackson, S.P., and Durocher, D. (2013). Regulation of DNA damage responses by ubiquitin and SUMO. Mol. Cell 49, 795–807.

Jiang, J., Maes, E.G., Taylor, A.B., Wang, L., Hinck, A.P., Lafer, E.M., and Sousa, R. (2007). Structural basis of J cochaperone binding and regulation of Hsp70. Mol. Cell 28, 422–433.

Kessler, B.M. (2013). Ubiquitin - omics reveals novel networks and associations with human disease. Curr. Opin. Chem. Biol. 17, 59–65.

Keuss, M.J., Hjerpe, R., Hsia, O., Gourlay, R., Burchmore, R., Trost, M., and Kurz, T. (2019). Unanchored tri NEDD8 inhibits PARP 1 to protect from oxidative stress induced cell death. EMBO J. e100024.

Kityk, R., Kopp, J., Sinning, I., and Mayer, M.P. (2012). Structure and dynamics of the ATP-bound open conformation of Hsp70 chaperones. Mol. Cell 48, 863–874.

Larance, M., Bailly, A.P., Pourkarimi, E., Hay, R.T., Buchanan, G., Coulthurst, S., Xirodimas, D.P., Gartner, A., and Lamond, A.I. (2011). Stable-isotope labeling with amino acids in nematodes. Nat. Methods 8, 849–851.

Leidecker, O., Matic, I., Mahata, B., Pion, E., and Xirodimas, D.P. (2012). The ubiquitin E1 enzyme Ube1 mediates NEDD8 activation under diverse stress conditions. Cell Cycle Georget. Tex 11, 1142–1150.

Lettre, G., and Hengartner, M.O. (2006). Developmental apoptosis in C. elegans: a complex CEDnario. Nat. Rev. Mol. Cell Biol. 7, 97–108.

Leu, J.I.-J., Pimkina, J., Frank, A., Murphy, M.E., and George, D.L. (2009). A small molecule inhibitor of inducible heat shock protein 70. Mol. Cell 36, 15–27.

Li, T., Guan, J., Huang, Z., Hu, X., and Zheng, X. (2014). RNF168-mediated H2A neddylation antagonizes ubiquitylation of H2A and regulates DNA damage repair. J. Cell Sci. 127, 2238–2248.

Liu, G., and Xirodimas, D.P. (2010). NUB1 promotes cytoplasmic localization of p53 through cooperation of the NEDD8 and ubiquitin pathways. Oncogene 29, 2252–2261.

Ma, T., Chen, Y., Zhang, F., Yang, C.-Y., Wang, S., and Yu, X. (2013). RNF111-Dependent Neddylation Activates DNA Damage-Induced Ubiquitination. Mol. Cell 49, 897–907.

Mahata, B., Sundqvist, A., and Xirodimas, D.P. (2012). Recruitment of RPL11 at promoter sites of p53-regulated genes upon nucleolar stress through NEDD8 and in an Mdm2-dependent manner. Oncogene 31, 3060–3071.

Martínez-Chantar, M.L., Vázquez-Chantada, M., Ariz, U., Martínez, N., Varela, M., Luka, Z., Capdevila, A., Rodríguez, J., Aransay, A.M., Matthiesen, R., et al. (2008). Loss of the glycine N-methyltransferase gene leads to steatosis and hepatocellular carcinoma in mice. Hepatol. Baltim. Md 47, 1191–1199.

Mayer, M.P., and Bukau, B. (2005). Hsp70 chaperones: Cellular functions and molecular mechanism. Cell. Mol. Life Sci. 62, 670–684.

Mergner, J., Heinzlmeir, S., Kuster, B., and Schwechheimer, C. (2015). DENEDDYLASE1 Deconjugates NEDD8 from Non-Cullin Protein Substrates in Arabidopsis thaliana. Plant Cell 27, 741–753.

Ordureau, A., Münch, C., and Harper, J.W. (2015). Quantifying ubiquitin signaling. Mol. Cell 58, 660–676.

Ouyang, H., Ali, Y.O., Ravichandran, M., Dong, A., Qiu, W., MacKenzie, F., Dhe-Paganon, S., Arrowsmith, C.H., and Zhai, R.G. (2012). Protein Aggregates Are Recruited to Aggresome by Histone Deacetylase 6 via Unanchored Ubiquitin C Termini. J. Biol. Chem. 287, 2317–2327.

Palleros, D.R., Welch, W.J., and Fink, A.L. (1991). Interaction of hsp70 with unfolded proteins: effects of temperature and nucleotides on the kinetics of binding. Proc. Natl. Acad. Sci. U. S. A. 88, 5719–5723.

Popovic, D., Vucic, D., and Dikic, I. (2014). Ubiquitination in disease pathogenesis and treatment. Nat. Med. 20, 1242–1253.

Qi, S., Pang, Y., Hu, Q., Liu, Q., Li, H., Zhou, Y., He, T., Liang, Q., Liu, Y., Yuan, X., et al. (2010). Crystal structure of the Caenorhabditis elegans apoptosome reveals an octameric assembly of CED-4. Cell 141, 446–457.

Rodriguez, J., and Lazebnik, Y. (1999). Caspase-9 and APAF-1 form an active holoenzyme. Genes Dev. 13, 3179–3184.

Saleh, A., Srinivasula, S.M., Balkir, L., Robbins, P.D., and Alnemri, E.S. (2000). Negative regulation of the Apaf-1 apoptosome by Hsp70 Nat. Cell Biol. 2, 476–483.

Shen, L., Liu, H., Dong, C., Xirodimas, D., Naismith, J.H., and Hay, R.T. (2005). Structural basis of NEDD8 ubiquitin discrimination by the deNEDDylating enzyme NEDP1. EMBO J. 24, 1341–1351.

Sundqvist, A., Liu, G., Mirsaliotis, A., and Xirodimas, D.P. (2009). Regulation of nucleolar signalling to p53 through NEDDylation of L11. EMBO Rep. 10, 1132–1139.

Tatham, M.H., Rodriguez, M.S., Xirodimas, D.P., and Hay, R.T. (2009). Detection of protein SUMOylation in vivo. Nat. Protoc. 4, 1363–1371.

Watson, I.R., Li, B.K., Roche, O., Blanch, A., Ohh, M., and Irwin, M.S. (2010). Chemotherapy induces NEDP1-mediated destabilization of MDM2. Oncogene 29, 297–304.

Williamson, A., Werner, A., and Rape, M. (2013). The Colossus of ubiquitylation: decrypting a cellular code. Mol. Cell 49, 591–600.

Wolter, K.G., Hsu, Y.T., Smith, C.L., Nechushtan, A., Xi, X.G., and Youle, R.J. (1997). Movement of Bax from the cytosol to mitochondria during apoptosis. J. Cell Biol. 139, 1281– 1292.

Xirodimas, D., Saville, M.K., Edling, C., Lane, D.P., and Laín, S. (2001). Different effects of p14ARF on the levels of ubiquitinated p53 and Mdm2 in vivo. Oncogene 20, 4972–4983.

Xirodimas, D.P., Sundqvist, A., Nakamura, A., Shen, L., Botting, C., and Hay, R.T. (2008). Ribosomal proteins are targets for the NEDD8 pathway. EMBO Rep. 9, 280–286.

Xu, G., Paige, J.S., and Jaffrey, S.R. (2010). Global analysis of lysine ubiquitination by ubiquitin remnant immunoaffinity profiling. Nat. Biotechnol. 28, 868–873.

Young, J.C. (2010). Mechanisms of the Hsp70 chaperone systemThis paper is one of a selection of papers published in this special issue entitled “Canadian Society of Biochemistry, Molecular & Cellular Biology 52nd Annual Meeting — Protein Folding: Principles and Diseases” and has undergone the Journal’s usual peer review process. Biochem. Cell Biol. 88, 291–300.

Zeng, W., Sun, L., Jiang, X., Chen, X., Hou, F., Adhikari, A., Xu, M., and Chen, Z.J. (2010). Reconstitution of the RIG-I Pathway Reveals a Signaling Role of Unanchored Polyubiquitin Chains in Innate Immunity. Cell 141, 315–330.

Zhao, Y., Brickner, J.R., Majid, M.C., and Mosammaparast, N. (2014). Crosstalk between ubiquitin and other post-translational modifications on chromatin during double-strand break repair. Trends Cell Biol. 24, 426–434.

Zhuravleva, A., Clerico, E.M., and Gierasch, L.M. (2012). An interdomain energetic tug-of-war creates the allosterically active state in Hsp70 molecular chaperones. Cell 151, 1296–1307.

