## Supplemental information for "The NEDD8 cycle controlled by NEDP1 upon DNA damage is a regulatory module of the HSP70 ATPase activity"

Bailly et al.

### Supplemental information

#### Experimental models

##### Worm strain and culture conditions

Worms were maintained at 20°C on NGM agar plates according to standard procedure unless otherwise indicated (Brenner, 1974). The following mutations and transgenes used in this study were: *ced-1(e1935), ced-3(n717), ced-9(n1653ts), cep-1(lg12501), gla-3(op216), glp-4(bn2ts), hpr-17(tm1579), lig-4(ok716), opls219* (CED-4::GFP), *orn-1(tm5454), ulp-3(tm1287).* The *ulp-3 (tm1287)* and *orn-1(tm5454)* deletion alleles were obtained from the Japanese Consortium (NBRP) and back-crossed five times against our N2 wild type genetic background. Double and triple mutants generated in this study will be available at the CGC and/or available on request. Apoptosis, cell cycle arrest assays and genotoxic sensitivity testing were performed as previously described (Bailly et al., 2010; Bailly and Gartner, 2013).

##### Cell culture

All cell lines were originally obtained from the ATCC bioresource. Cell lines were maintained in DMEM in 10% FCS and standard antibiotics, in 5% CO2 and 37$^{o}$C and regularly tested for mycoplasma contamination. Cell lines have not been authenticated.

##### Mouse model systems

Animal experiments were performed according to the Spanish Guide for Care and Use of laboratory animals and the European Research Council animal care and use guidelines. Protocols were approved by the CIC bioGUNE Animal Care and Use Committee and the local authority (Diputación de Bizkaia). In the present study, animals were between 5 to 12-month-old males. Animals were bred and housed in the animal unit of CIC bioGUNE (AAALAC-accredited facility) with controlled conditions of temperature (22ºC) and humidity, in a 12 h light/dark cycle with *ad libitum* access to food and water.

#### Materials and Methods

##### Worm protein extraction and fractionation

For Western Blots, worm proteins were extracted from synchronised adult worms in 50mM Hepes-NaOH pH8.0, Urea 9M, SDS 2% supplemented with zirconium 0.7mm beads (BioSpec Product) and disrupted in a bead beater (FastPrep, 3 cycles of 6m/sec, 20sec each). Debris were removed by quick centrifugation cycles (1000g, 1min, 3 times) and soluble proteins were then separated by a 12000g, 30min centrifugation. Supernatant protein concentration was measured by BCA following manufacturer's instructions (Life technologies). 2xSDS Laemmli buffer was added before boiling samples for 5min.

##### Stable Isotope Labelling with Amino acid in Nematodes (SILAC)

##### Worm Isotope labelling

To minimize Arginine-to-Proline conversion we obtained a deletion allele from the Japanese NBRP consortium that disrupts the *orn-1* gene *(tm5454)*. We back-crossed the *orn-1(tm5454)* deletion five times against our N2 wild type background then against *ulp-3(tm1287).* For isotopic labelling, SILAC bacteria (SLE1) were labelled as in (Larance et al., 2011). Synchronised worms were grown on nitrogen free nematode growth medium (NGM-N) as described previously (Larance et al., 2011) seeded with SLE1 bacteria labelled with Light and Heavy Isotopes (Eurisotope). Larvae stage 1 synchronised worms were grown until late adult stage and synchronised again by bleaching before growing them on 9cm NGM-N plates seeded with SILAC bacteria (SLE1). Approximately 10 plates of 4000 worms each were used to grow synchronised and labelled worms until late L4-young adult stage that were irradiated with 90Gy of IR using a Blood-Xrad apparatus. Labelled worms proteins were extracted in 50mM Hepes-NaOH pH8.0, Urea 9M using a bead beater and zirconium 0.7mm beads following the same procedure as described in the protein extraction section.

##### Mass spectrometry

Di-Glycine motif peptide identification was performed following UbiScan protocols and instructions (Cell Signaling Technology) using Ubiquitin Branch Motif Antibody (K-ε-GG) ♯3925. Peptides were loaded onto 10cm x 75µm PicoFrit Capillary column packed with Magic C18 AQ reversed-phase resin. The column was developed with a 90min linear gradient of acetonitrile in 0.125% formic acid delivered at 280nL/min. MS parameters settings: MS run time 96min, MS1 Scan Range (300.0-1500.00), Top 20 MS/MS (Min Signal 500, isolation width 2.0, normalized coll. energy 35.0, activation-Q 0.250, activation time 20.0, lock mass 371.101237, charge state rejection enabled, charge state 1+ rejected, dynamic exclusion enabled, repeat count 1, repeat duration 35.0, exclusion list size 500, exclusion duration 40.0, exclusion mass width relative to mass, exclusion mass width 10 ppm). MS/MS spectra were evaluated using SEQUEST 3G and the SORCERER 2 platform from Sage-N Research (v4.0, Milpitas CA). Searches were performed against the most recent update of the NCBI *C. elegans* database with mass accuracy of +/- 50ppm for precursor ions and 1Da for product ions. Results were filtered with mass accuracy of +/- 5 ppm on precursor ions and presence of the intended motif (K-ε-GG). LTQ-Orbitrap Velos was used (Thermoscientific). A 5% default false positive rate was used to filter the SORCERER results.

**Worm Live Imaging Preparation**

For FRAP-based acquisitions, worms were individually picked on an unseeded NGM plate to wash off bacteria and transferred on a glass slide in a 10µl drop of egg buffer (118mM NaCl, 48mM KCl, 2mM $CaCl_2$*2$H_2O$, 2mM $MgCl_2$*6$H_2O$, 25mM HEPES pH 7.3). Worm gonads were dissected with a 23G syringe and immediately covered with a cover slip sealed with nail polish. Acquisitions were performed immediately after (less than 5min).

**Fluorescence Recovery After Photobleaching (FRAP)**

FRAP analysis was performed on a confocal microscope Leica Sp5, objective Leica 63x/1.4 oil Apo. GFP was photo-bleached by "FRAP wizard" function of Leica LAS software. The region of one nucleus was photo-bleached with maximal 488nm laser to obtain more than 70% of photo-bleaching and recovery with 5% of laser power. Fluorescence intensities were measured by five acquisitions in pre-bleached and twenty acquisitions every 0.34sec (post–bleached 1) and every 10sec nine times during the recovery (post-bleached 2). Intensities data of bleach, background and non-bleached nucleus regions (total intensity) were extracted from the Leica software and analysed with excel to calculate fluorescent protein mobility. The fraction of the fluorescent protein that was mobile was calculated by comparing the fluorescence intensities of non-bleached and bleached nucleus.

***C. elegans* RNAi**

We adapted the RNAi bacterial feeding method from (Larance et al., 2011*)*. Briefly, *E. coli* HT115 RNAi bacteria were grown in M9 minimal supplemented with glucose 0.4% (wt/vol), 2mM MgSO4, 0.1mM $CaCl_2$, thiamine 0.01% (wt/vol) and 100µg/ml ampicillin. A single bacteria colony streaked on LB plate was used to start the culture at 37°C under agitation until OD600nm = 1. Double-stranded RNA expression was induced by adding IPTG to a final concentration of 0.5mM for 3hrs. Bacteria culture was seeded on NGM plates complemented with ampicillin. Worms were placed on NGM plates seeded with RNAi bacteria at late L1 stage for *ulp-3* RNAi and L3 stage for *ned-8* RNAi. Late L4 to young adult stage worms were analysed. We found that growing HT115 bacteria in minimal medium greatly increases RNAi efficiency even for loci reported to be problematic to achieve appropriate depletion level.

For *ulp-3* and *ned-8* RNAi, the following oligos were used to clone cDNA into the L4440 vector:

For *ulp-3*:

5' to 3': TATATATAGCGGCCGCATGTCAGTTCCCCCGG

5' to 3': TATATATAGCGGCCGCTTATTTCGCTTCAAAATTAACAATC

For *ned-8*:

5' to 3': AGGCGGCCGCTTAAAATCCTCCGCGG

5' to 3': AAGGCGGCCGCATGCTCATCAAAGTTA

**qRT-PCR experiments**

Gravid adult worms were collected from the plates and washed three times to remove bacteria with PBS before introduction in Trizol and frozen at -80°C. Total RNA was extracted by Trizol/chloroform followed by RNeasy® Kit (Qiagen) and cDNA synthesis performed on 300ng of RNA with QuantiTech® Reverse Transcription Kit (Qiagen). qRT-PCR reactions were performed in Roche LC480 qPCR in triplicates, and cDNA was diluted in Mesa Green qPCR Master Mix (Eurogentec) with specific primers. The mean values for the levels of transcript of interest (*egl-1*) were normalised with internal level of control transcript (tubulin *tbg-1*) relatively stably expressed between all experiments. For data analysis, maximum quantitative point method was applied and induction was calculated with the equation:

Fold Induction= 2^(($^{\Delta Cp(cDNAegl\text{-}1\text{-}cDNAtbg\text{-})}$control)-($^{\Delta Cp(cDNAegl\text{-}1\text{-}cDNAtbg\text{-})}$treated)). Data are represented as mean +/- SEM.

**ULP-3 antibody production**

Full length ULP-3 was amplified from cDNA derived from adult worms using the same protocol as used in qRT-PCR experiments section using the following oligo:

5'-TATATATAGGCGCGCCATGTCAGTTCCCCCGGATTTCCGC (forward)

5'-GAAGCGAAATAAGGCCGGCCTATATATA (reverse). AscI was introduced at the 5' of the start codon and FseI at the 3' of the stop codon and cloned into appropriately modified pQE-80L ($His_6$) and pMAL-c2 (MBP). Protein expression of MBP-ULP-3 and $His_6$-ULP-3 was performed in BL21(DE3) CodonPlus bacteria grown and induced at 37°C for 2hrs with 0.5mM IPTG. MBP-ULP-3 was purified according to manufacturer standard protocols for amylose resin (New England Biolabs). $His_6$-ULP-3 was purified on Cobalt Talon beads (Clontech) following manufacturer instructions for soluble protein and finally dialysed three times against 50mM Tris-HCl pH7.5, 150mM NaCl, 10% glycerol, 0.5mM DTT. MBP-ULP-3 was used to immunize rabbits (BioGenes GmbH). For affinity purification of the antibodies, $His_6$-ULP-3 was covalently coupled to AffiGel-15 resin (BioRad) and used to purify final bleeds serum.

**ULP-3 and NEDD8 site-directed mutagenesis**

A catalytically inactive mutant of MBP-ULP-3 (C167A) was generated using the following oligos:

5'-GCATCCGATGCTGGACTCC, 5'-GGAGTCCAGCATCGGATGC. The hydrophobic patch NEDD8 mutant (I44A) was generated using the following oligos:

5'-GCAGAGGCTCGCCTACAGTGGCAAG, 5' -CTTGCCACTGTAGGCGAGCCTCTGC

Sequences were verified by automated sequencing.

**Reagents**

Aldolase and thyroglobulin were purchased from SIGMA and BioRad Laboratories respectively. The Flag-L11 and NEDP1 constructs were described previously (Sundqvist et al., 2009). ULP-3 was cloned in pcDNA3 vector as Flag-tagged (N-terminus). The catalytic cysteine mutant was created by side directed mutagenesis and sequences were confirmed by automated sequencing. Reagents for Annexin V staining were from BD Biosciences, MLN4924 from Active Biochem. Prestained Protein Ladders, from Thermo Scientific and BioRad. All protein samples were analysed in 4-12% precast SDS-PAGE-Invitrogen. Antibodies: Mouse β-actin (Ab-1, Calbiochem), rabbit APAF-1 (E38 for IPs, Abcam), mouse APAF1 (input, BD Biosciences, 611364), mouse cullin 1 (Zymed, 32-2400), rabbit H2Ax (ab11175, Abcam), rabbit gH2Ax (05-636, Millipore), rabbit H2A (ab13923, Abcam), mouse HSP70 (ab47455, Abcam), rabbit NEDD8 (Y297, GeneTex), mouse p53 (DO-1, in house), sheep NEDP1 (in house), rabbit ULP-3 (in house), mouse Tubulin (SIGMA, T9026), mouse Flag and anti-Flag M2 affinity gel (SIGMA, A2220), rabbit ubiquitin (DAKO, z0458), mouse GAPDH (Santa Cruz, 6C5, sc-32233).

**Real time PCR-Human cells/Mice**

RNA from cells in 6cm dishes was isolated with the Promega SV Total RNA Isolation kit. cDNA synthesis was performed using the Invitrogen SuperScript III First-Strand Synthesis SuperMix using 300ng of RNA. Real-time PCR was performed as described in in (Mahata et al., 2012) an ABI 7500 system using ABI PCR master mix. The TaqMan primers for p21 can be found in (Mahata et al., 2012), whereas the NEDP1 primers were purchased from ABI. All experiments were performed at least 3 times and error bars represent +/- SEM. RNA isolation from liver tissue was isolated using TRIZOL/Chloroform and RNA quantification was determined in the Nanodrop ND-100 spectrophotometer (ThermoFischer Scientifict, USA). 2μg of isolated RNA were treated with DNase I and cDNA was synthesized with M-MLV reverse transcriptase in the presence of random primers and RNaseOUT. Real time PCR was performed with 1.5μl of cDNA (diluted 1:10), SYBR Select Master Mix. Primers for monitoring nedp1 (senp8) mRNA expression in mice: F: 5'ATCCCTGCTCAATACAAGATGGA, R: 5' CAAACCCAATAATGTGGTCGTTG. Data were normalized against gapdh expression, F: 5'GGATGCAGGGATGATGTTC and R: 5'TGCACCACCAACTGCTTAG.

**Tissue culture and siRNA transfections**

All cell lines were originally obtained from the ATCC bioresourse. Cell lines were maintained in DMEM in 10% FCS and standard antibiotics, in 5% CO2 and 37$^{o}$C and regularly tested for mycoplasma contamination. For siRNA transfections 5nM of oligos were transfected with RNAiMAX lipofectamine (Life Technologies). All siRNAs are from Dharmacon as ON-TARGET-Plus pools of 4 oligos.

**Stable cell lines**

U2OS cells stably expressing wild type $His_6$-NEDD8 or $His_6$-NEDD8 K11/48R mutant was performed as described in (Liu et al., 2010), using lentivirus with the respective constructs. Cells were selected and maintained in puromycin (2.5μg/ml). This method provides clones expressing low levels of ectopic NEDD8 similar to the endogenous levels (Liu et al., 2010). Several assays (treatment with MLN4924, or proteotoxic stress) confirmed that the ectopic NEDD8 responds similarly to the endogenous NEDD8. Parental or H6 NEDP1 KO U2OS cells were transfected with either empty or 3xFlag-HSP70 pcDNA3 vectors. Cells were selected with G418 (1mg/ml) for 14 days before a pool of cells stably expressing 3xFlag-HSP70 was acquired.

**Isolation of $His_6$-NEDDylated/ubiquitinated proteins proteins**

H1299 cells in 10cm dishes were transfected with 1μg of $His_6$-NEDD8 or $His_6$-ubiquitin, 2μg of Flag-L11, 3μg of Flag-ULP-3 or NEDP1 with Fugene. 48hrs post transfection $Ni^{2+}$-pull downs and total cell extracts (input 2xSDS Laemmli buffer) were prepared as described in ( Xirodimas et al., 2001, Tatham et al. 2009).

**Caspase 3/7 assay-Annexin V staining**

U2OS cells were transfected in 6 well plates. Next day cells were trypsinised and seeded in 24well plates (7000 cells/well) in duplicates. 24hrs later cells were treated as indicated and one set of cells was used to measure cell survival (CellTiter-Glo® Luminescent Cell Viability Assay, Promega) and the other caspase 3/7 activity (Caspase-Glo 3/7 Assay, Promega). The ratio between caspase 3/7 activity over survival was measured and represented as fold change to the control untreated cells. For Annexin V staining 1x10cm 80% confluent U2OS cells was used for each condition. Cells were collected by careful trypsinisation (floating cells were also included), washed in PBS, before resuspended in binding buffer, $1x10^6$ cells/ml (10mM HEPES/NaOH, pH 7.4, 150mM NaCl, 5mM KCl, 5mM $MgCl_2$, 1.8mM $CaCl_2$) and FITC-Annexin V was added at 1:100 dilution. Samples were kept at RT in the dark for 20min before 7-ADD viability dye (5μg/sample, late apoptotic cells) was added and samples were analysed by flow cytometry. For both assays data represent the average of 3 different experiments +/- standard deviation.

**CRISPR/Cas9 deletion of NEDP1**

The U6gRNA-Cas9-2A-GFP plasmid for NEDP1 was purchased from SIGMA. 2 plasmids with different guide sequences were obtained (#20, #22). U2OS cells were transfected in 10cm dishes with the plasmids and 24hrs later cells were collected and GFP positive cells were selected and separated in 96-well plates as single cells by flow cytometry. Several clones were screened for NEDP1 and NEDD8 expression. Total DNA from the used clone in this study (H6) was used to amplify the NEDP1 locus by PCR and sequencing of the PCR product confirmed the deletion in NEDP1 gene (305-314bp).

**Subcellular fractionation**

2x10cm dishes of 80% confluent parental or H6 U2OS cells were used for fractionation. Cells were washed and collected in 1ml PBS and 100μl of the sample were pelleted at 13000rpm for 1min and lysed with 100μl of 2xSDS Laemmli buffer (input). The remaining 900μl were pelleted at 10800rpm for 20s and resuspended in 300μl buffer A (10mM HEPES-KOH pH 8.0, 10mM KCl, 1.5mM $MgCl_2$) with protease inhibitor (complete EDTA-free, Roche) and 10mM iodoacetamide. Cells were lysed by adding Triton-X-100 at a final concentration of 0.1% for 1min at 4°C, then centrifuged for 5min at 1300g, 4°C. The supernatant (cytoplasmic fraction) was mixed with equal volume of 2xSDS Laemmli buffer. The pellet (nuclear fraction) was washed 3x with buffer A, then resuspended with 300μl of buffer B (20mM HEPES-KOH pH 8.0, 300mM NaCl, 2mM EDTA and 1%NP40) and incubated 30min on ice. Lysates were centrifuged at 10800rpm for 20s. The supernatant (nucleoplasmic fraction) was mixed with equal volume of 2xSDS Laemmli buffer. The pellet was washed 3 times with buffer B then resuspended in 600μl of 2xSDS Laemmli buffer (pellet fraction).

**Immunoprecipitations**

For each condition 1x10cm dish of 80% confluent U2OS cells was used. Cells were washed 2x with PBS before collected and lysed in 50mM Tris, pH 7.4, 100mM KCl, 1% NP-40, 2mM EDTA, protease inhibitors (complete EDTA-free, Roche) for 15min on ice. Lysates were cleared by centrifugation at 13000rpm for 15min at 4°C and extracts of equal amount of total protein were used for immunoprecipitations overnight at 4°C with 2μg of APAF1 antibody and 20μl of protein G beads (prewashed with lysis buffer). Next day beads were washed 2x with 500μl of lysis buffer (centrifuged at 5000rpm at 4°C in each wash), resuspended in 100μl of 2xSDS Laemmli buffer and boiled for 5min. Immunoprecipitates were analysed by western blotting. For Flag pull downs 20μl of pre-washed Flag beads were used.

**In vitro NEDD8 processing assay**

Protein induction in bacteria was performed with 0.5mM IPTG at 16°C overnight and induced proteins were purified as MBP fusions using standard methods. NEDP1 was purified as $His_6$-GST fusion before cleavage with thrombin as described in (Shen et al., 2005). 600ng of MBP-NEDD8-ubiquitin fusion and 200ng of ULP-3/NEDP1 proteins were incubated for the indicated periods of time at 37°C in 10μl of 50mM Tris, pH 7.5, 100mM NaCl, 5mM $MgCl_2$, 1mM DTT. Reactions were terminated with the addition of 10μl of 2xSDS

Laemmli buffer, samples were boiled for 5min, separated on a 4-12% precast SDS-PAGE and coomassie blue stained.

**Sucrose gradient fractionation**

4x10cm dishes of 80% confluent cells were washed 2x with PBS before collected and lysed in 50mM Tris, pH 7.4, 100mM KCl, 1% NP-40, 2mM EDTA, protease inhibitors (Roche tablets) for 15min on ice. After centrifugation at 14000rpm for 15min at 4$^o$C, equal amount of extracts were loaded on 10-50% sucrose gradient. Samples were centrifuged for 24hrs at 36000rpm at 4$^o$C in SW41 Ti swing out rotor. 1ml fractions were collected using the Brandel density gradient fractionation system with upward displacement. Proteins were TCA precipitated, pellets washed 3x with cold acetone resuspended in 100μl of 2xSDS Laemmli buffer, boiled for 5min and samples were analysed by western blotting. On a separate experiment samples were spiked with aldolase (158kDa) and thyroglobulin (690kDa) as controls for the fractionation.

**Plasmids**

NEDD8, HSPA1A, DNAJB4, DNAJA1, NEDP1 cDNA (Montpellier Genomic Collection) were cloned into pET15b plasmids for $His_6$-tagged constructs. For GST tagged versions, HSP70 constructs (residues 1-641, 1-383, 1-402, 1-135, 190-402) were cloned into pGEX-2TK. For point mutants, the DpnI site directed mutagenesis method was followed.

**Purification of recombinant $His_6$-NEDD8 constructs.**

***Critical***: All *in vitro* experiments were performed with freshly produced/purified $His_6$-NEDD8 proteins each time, with no prior freezing. Constructs were expressed in BL21 codon plus bacteria grown and induced at 37°C for 2hrs with 0.5mM IPTG and lysed immediately in 50mM Tris-HCl pH7.5, 0.5M NaCl, 10% Glycerol, 10mM Imidazole, 0.2% Tween-20 and 5mM β-mercaptoethanol at 4°C for 30min then sonicated (3x 45% amplitude, 30 sec) and centrifuged at 20000g for 30min at 4°C. Supernatant was incubated with 300μl of pre-washed nickel beads for at least 4hrs at 4°C. Beads were extensively washed in lysis buffer and finally washed in 50mM Tris-HCl pH7.5, 150mM NaCl, 5mM β-mercaptoethanol. Material was then eluted in 20mM Tris-HCl pH7.5, 150mM NaCl, 0.5M Imidazole, 5mM β-mercaptoethanol. After analysis by PAGE followed by Coomassie Blue staining, fractions were dialysed in 20mM Tris-HCl pH7.5, 150mM NaCl, 0.5mM DTT and stored at 4°C for subsequent *in vitro* analysis.

**Purification of recombinant GST HSP70 constructs.**

Constructs were expressed in BL21 codon plus bacteria grown and induced at 37°C for 2hrs with 0.5mM IPTG. Cells were harvested and lysed in 50mM Tris-HCl pH7.5, 50mM NaCl, 0.5mM DTT and sonicated (amplitude 50%, 3 bursts of 30 sec). After 20000g centrifugation supernatant was incubated with 100μl of pre-washed GST beads for at least 4hrs at 4°C. Beads were then extensively washed with 50mM Tris-HCl pH7.5, 150mM NaCl, 0.1% Triton X-100 with a final wash in 50mM Tris-HCl pH7.5, 150mM NaCl, 0.5mM DTT. Sequential elutions were performed with 50mM Glutathione pH7.5, 50mM Tris-HCl pH7.5, 150mM NaCl, 0.5mM DTT. Eluted fractions were analysed by SDS-PAGE stained with Quick Coomassie and best fractions were pooled and dialysed against 50mM Tris-HCl pH7.5, 150mM NaCl, 0.5mM DTT. Extracts were finally cleared by 30000g centrifugation.

**Measurements of ATPase activity of HSP70**

Bacterially expressed $His_6$-HSP70 ATPase domain (residue 1-402, unless otherwise indicated) was prepared (final concentration 2µM) in assay buffer (100mM Tris-HCl pH7.5, 20mM KCl, 6mM $MgCl_2$) in 50µl final volume. This construct contains the linker domain which stimulates the ATPase activity of Hsp70 (Jiang et al., 2007). BSA, NEDD8, Ubiquitin or HSP40 were added at the indicated concentration and incubated at 4°C for 2hrs to allow binding. The reaction was started by adding 0.5µl of 20mM ATP. After 2hrs of incubation at 37°C, 25µl of the reaction mixture were added to 75µl of green malachite reagent (SIGMA, ATPase activity assay kit) on a 96-well plate and incubated for 30min at room temperature. The absorbance was determined at 620nm with a Polarstar Omega apparatus (BMG). To correct for non-enzymatic hydrolysis of ATP, the absorbance of a sample lacking HSP70 but otherwise treated identically was subtracted.

**Isolation of poly-NEDD8 conjugates from cell extracts and ATPase activity measurement**

U2OS cell lines stably expressing $His_6$-NEDD8 and deleted for NEDP1 were incubated in lysis buffer (20mM Tris-HCl pH7.5, 0.5M NaCl, 10mM Imidazole, 0.2% Tween-20, 5mM β-mercaptoethanol) at 4°C for 30min then sonicated (30% amplitude, 5sec burst) and centrifuged at 20000g for 30min at 4°C. Supernatant was incubated with 50µl of pre-washed nickel beads for at least 4hrs at 4°C. Beads were extensively washed in high salt washing buffer (20mM Tris-HCl pH7.5, 1M NaCl, 10mM Imidazole, 0.2% Tween 20, 5mM β-mercaptoethanol) and finally washed three times with washing buffer without Tween-20. Material was then eluted in elution buffer (20mM Tris-HCl pH7.5, 0.5M NaCl, 500mM Imidazole, 5mM β-mercaptoethanol) twice in 100µl then dialysed against 100mM Tris-HCl pH7.5, 20mM KCl, 6mM $MgCl_2$ at least 10000 fold. For subsequent ATPase activity measurements, 20µl were incubated with or without 500ng of bacterially expressed $His_6$-NEDP1 at 37°C for 2hrs in 100µl final of 100mM Tris-HCl pH7.5, 20mM KCl, 6mM $MgCl_2$. Reactions were then transferred at 4°C and 2µM of HSP70 ATPase domain (1-402) were added and incubated for 2hrs to allow binding. Reactions were started by adding ATP at 200µM final concentration and incubated at 37°C for 2hrs. ATPase activity was measured as described above.

**In vitro binding assay**

1µg of bacterially expressed GST or various GST-HSP70 constructs were incubated with 300ng of bacterially expressed $His_6$-NEDD8 in 300µl binding buffer (50mM Tris-HCl pH7.5, 150mM NaCl, NP-40 1%, 1mM DTT) for 1hr at 4°C then 20µl of GST beads were added to each reaction for 1hr at 4°C under gentle rotation. Beads were washed 3x with binding buffer and 50µl of SDS loading buffer was added before boiling. Supernatants were analysed by western blotting.

**In vitro NEDDylation assay**

50µl reactions were set up including recombinant proteins: 300ng GST-NAE, 5µg Ubc12 and 10µg $His_6$-NEDD8 in 50mM Tris-HCl, pH 7.5, 2mM ATP, 5mM $MgCl_2$, 2mM DTT. Reactions were incubated at 37°C for 5hrs. $His_6$-NEDD8 conjugates were isolated with 10µl of pre-washed $Ni^{+2}$-agarose beads in 500µl of 50mM Tris-HCl, pH 7.5, 0.5M NaCl for 2hrs at room temperature. Beads were extensively washed with Tris-HCl, pH 7.5, 0.5M NaCl before elution (2x100µl) in Tris-HCl, pH 7.5, 0.5M NaCl, 250mM imidazole. Dialysis and ATPase activity measurements were performed as with the isolation of poly-NEDD8 conjugates from cell extracts (see above).

**GST pull-down from cell extracts**

Harvested cells were incubated in binding buffer (100mM Tris-HCl pH7.5, 20mM KCl, 6mM $MgCl_2$) at 4°C for 30min and sonicated (30% amplitude, 5sec burst) then centrifuged at 20000g for 30min at 4°C. Supernatant was incubated with 10µg of GST or GST-HSP70 constructs at 4°C at least 2hrs. Then 50µl of pre-cleared (binding buffer supplemented with 20µg of molecular grade BSA) GST beads were added and incubated for 1hr under gentle rotation. Beads were washed 3x with binding buffer supplemented with 0.5% NP-40 and 50µl of SDS loading buffer was added before boiling. Eluates were analysed by western blotting.

**Protein extraction from Liver Tissue**

Approximately, 50µg of liver tissue were homogenised by using Precellys 24 (Bertin Technologies) with 1ml of lysis buffer (1.6mM NaH2PO4, 8.4mM Na2HPO4, 0.1% Triton X-100, 0.1M NaCl, 0.1% SDS, 0.5% sodium azide) supplemented with protease and phosphatase inhibitor cocktail, 20mM iodoacetamide and 20mM N-Ethylmaleimide. Lysates were centrifuged (13000rpm, 30min, 4ºC) and total protein content in the supernatant was measured by Bradford. 20µg of protein was used for SDS-PAGE and western blot analysis.

Supplemental Figure 1

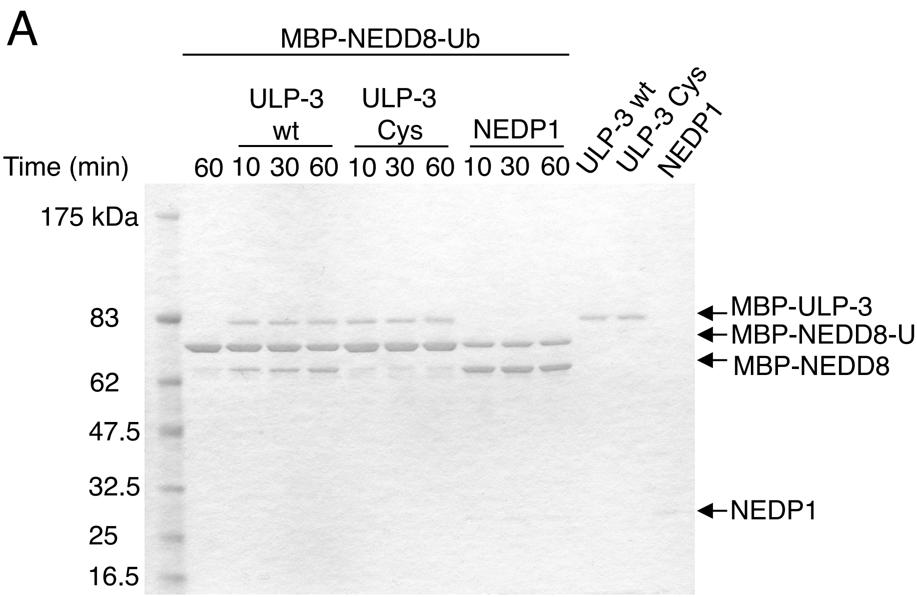


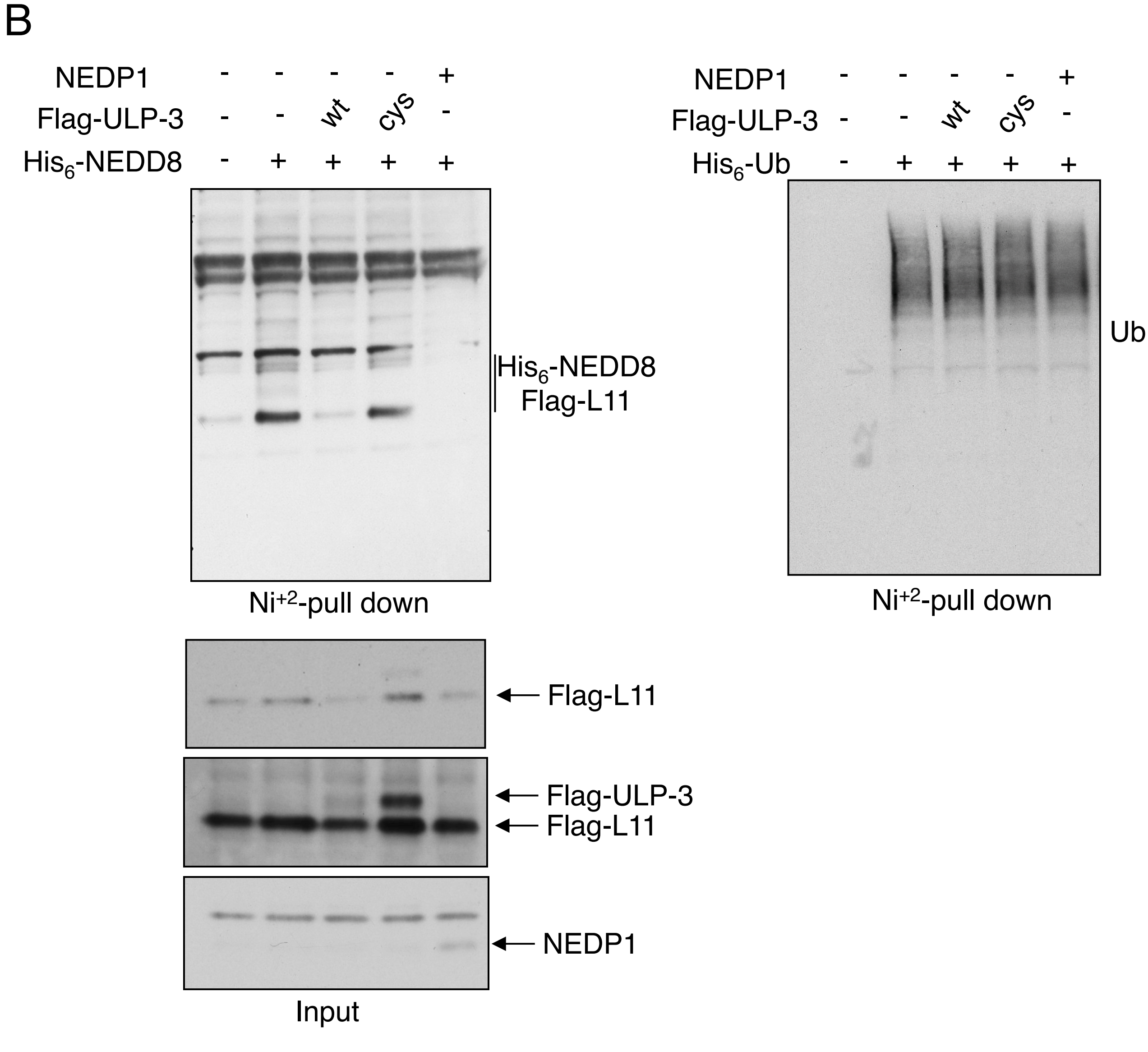

**Supplemental Figure 1**

**ULP-3 has NEDD8 processing and de-NEDDylating activity. A.** Bacterially expressed wild type MBP-ULP-3 or Cys catalytic mutant were tested *in vitro* for NEDD8 processing activity using the MBP-NEDD8-Ub fusion. As control wild type bacterially expressed NEDP1 was used. **B.** H1299 cells were transfected with the indicated constructs. In all samples Flag-L11 was also co-transfected. 48hrs post-transfection $His_6$-NEDD8 conjugates were isolated with Nickel beads and western blot analysis was performed with anti-Flag antibody. Total cell extracts (input) were analysed with the indicated antibodies. Similar experiments were performed using $His_6$-ubiquitin and after Nickel purification ubiquitin conjugates were detected with anti-ubiquitin antibody (right panel).

### Supplemental Figure 2

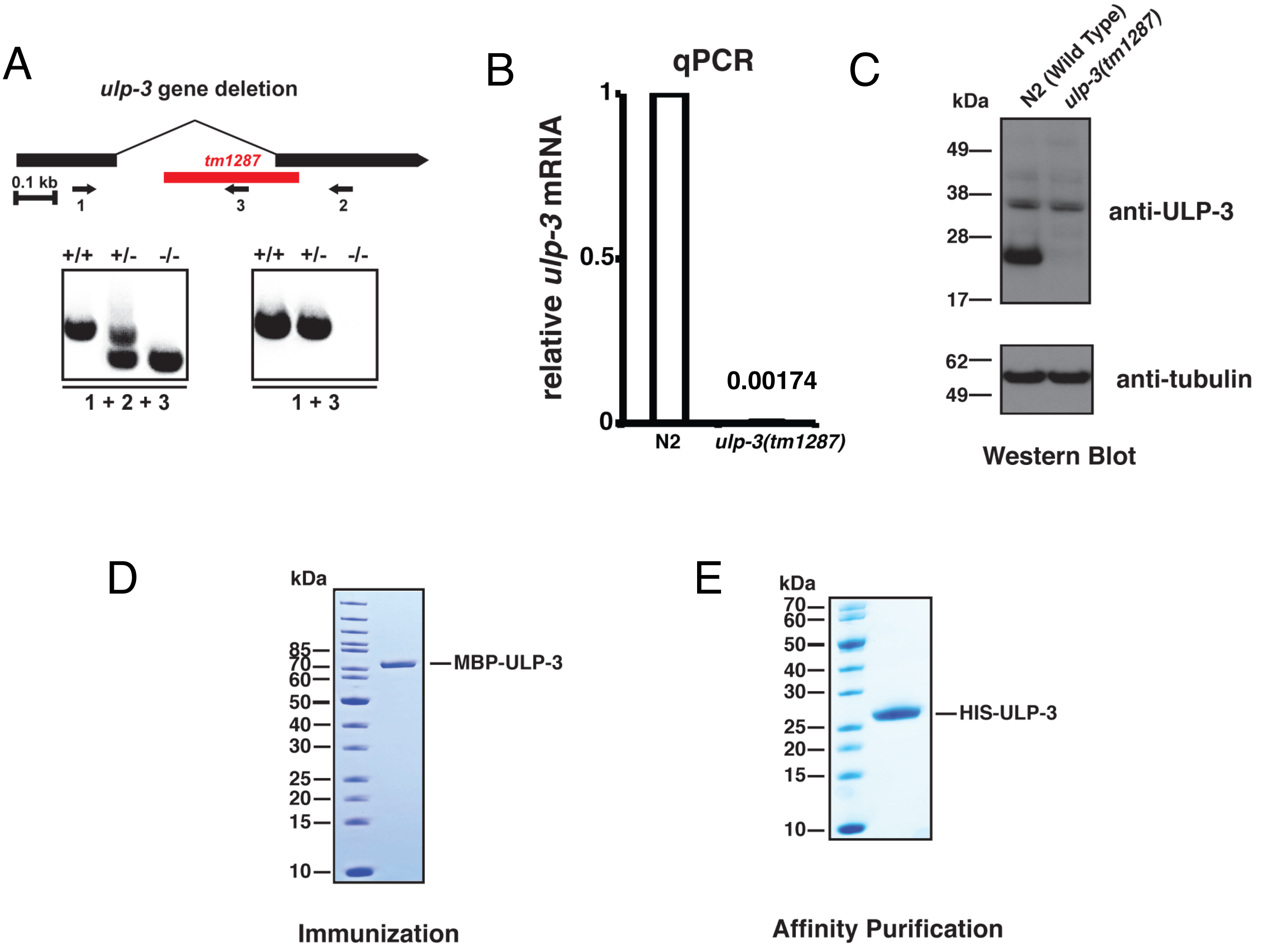


**Supplemental Figure 2**

**The *ulp-3(tm1287)* allele abolishes *ulp-3* gene expression. A.** The *tm1287* allele structure demonstrated by the PCR products generated by the indicated primers. +/+, +/- and -/- indicate the N2/N2, N2/*ulp-3* and *ulp-3/ulp-3* genotypes respectively. **B.** The gene disruption is demonstrated by the *ulp-3* mRNA level measured by real time PCR. Relative expression levels were determined using γ-tubulin as standard (*tbg-1*). Fold induction was calculated as previously described (Bailly et al., 2010). **C.** Wild type and *ulp-3(tm1287)* mutant worms were synchronised and grown under standard conditions. Protein extraction was performed as described in Methods and the ULP-3 protein levels analysed by Western blotting. α-tubulin was used as protein loading control. **D.** Recombinant MBP-ULP-3 protein, used for deNEDDylation *in vitro* assays and rabbit immunisation (Supplemental Figure 1A), was produced in BL21 cells and purified with amylose resin as described in Methods. Purified MBP-ULP-3 protein was analysed by SDS-PAGE followed by Coomassie Blue staining. **E.** Recombinant His-ULP-3 (used to purify ULP-3 antibodies) was produced in BL21 cells and purified on a Cobalt column then analysed by SDS-PAGE followed by Coomassie Blue staining.

### Supplemental Figure 3

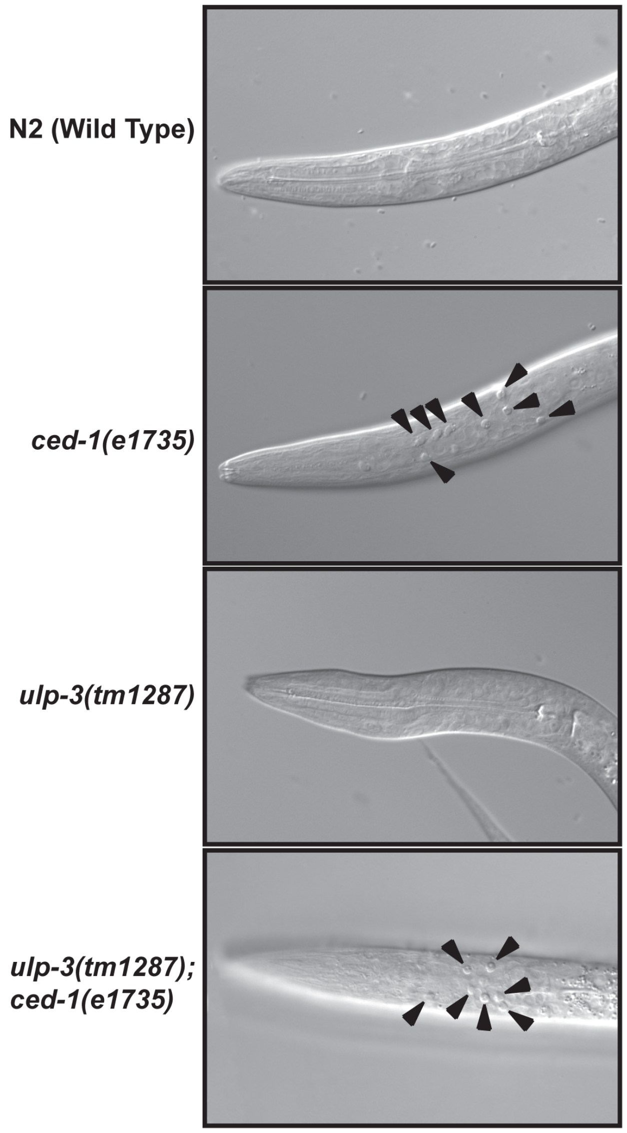


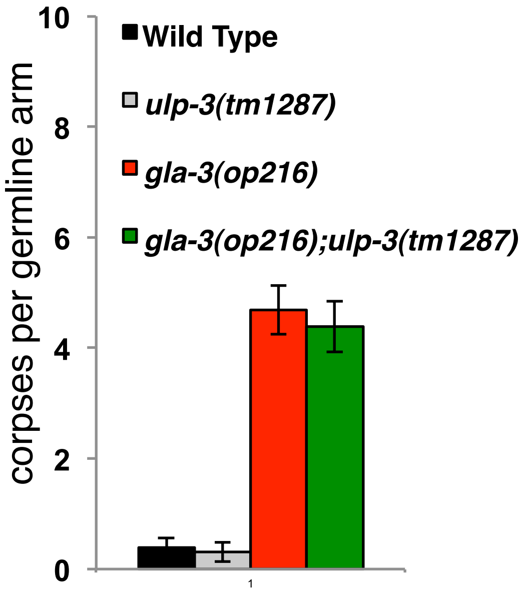


| Genotype | apoptotic corpses in the head |
|---|---|
| **Wild Type** | **0 ± 0** |
| ***ced-1(e1735)*** | **15.18 ± 0.65** |
| ***ulp-3(tm1287)*** | **0 ± 0** |
| ***ulp-3(tm1287); ced-1(e1735)*** | **15.09 ± 0.75** |

**Supplemental Figure 3**

**ULP-3 specifically affects the DNA damage-induced apoptotic pathway**. Upper panel: the apoptotic corpses that persist in the head until the L1 larvae stage in *ced-1(e1735)* animals were scored as in (Ellis et al., 1991). Representative DIC pictures of the indicated genotype are shown. Average number of corpses, +/- SEM is represented (*n=15*) in the lower panel. In contrast to wild type, the *gla-3(op216)* mutant displays an elevated number of germ cell apoptosis without DNA damage referred as physiological apoptosis level. The double mutant *ulp-3(tm1287); gla-3(op216)* does not show a significant decrease in the number of apoptotic cells suggesting that ULP-3 does not regulate physiological germ cell apoptosis (right panel).

Supplemental Figure 4

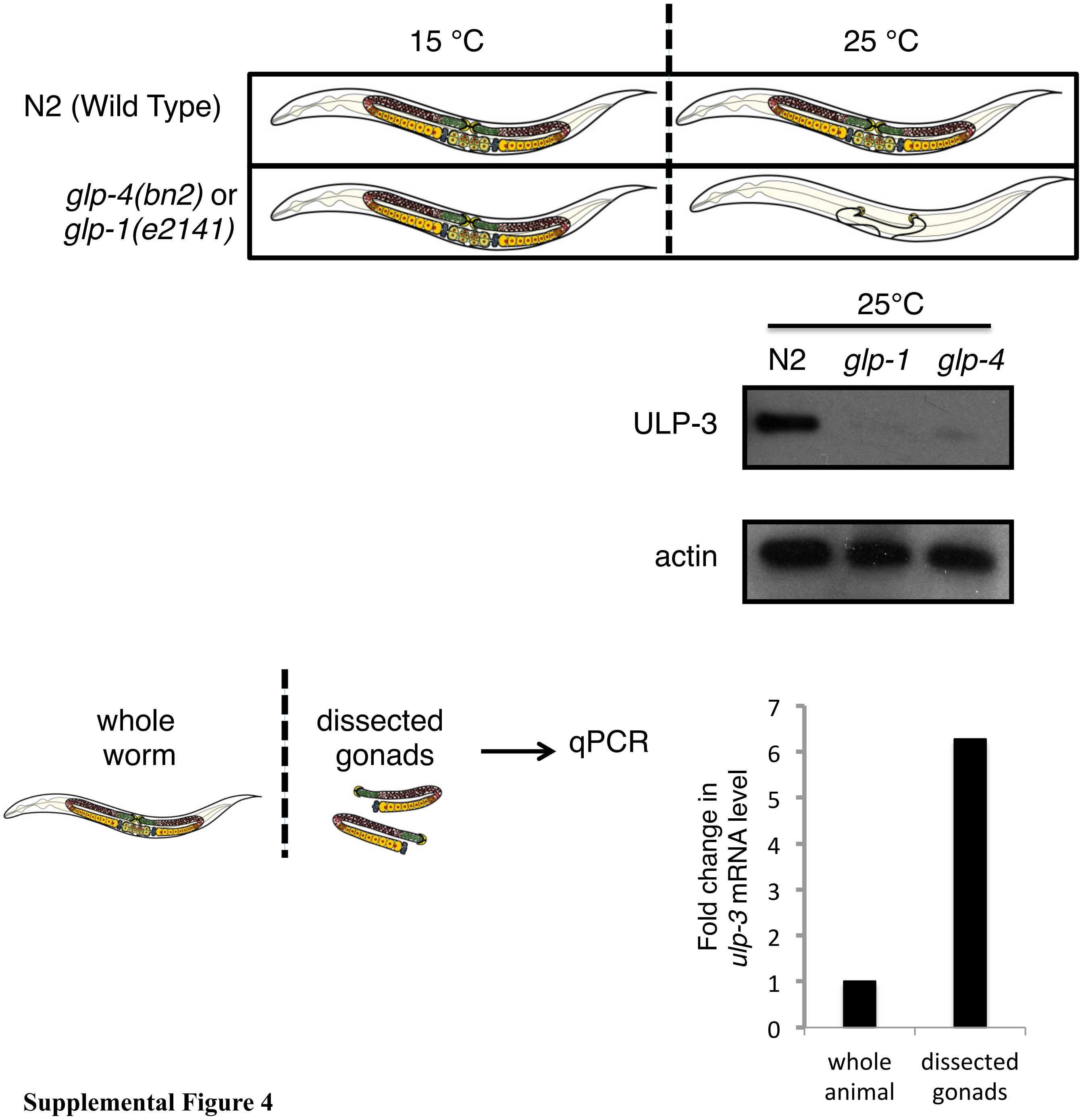


**Supplemental Figure 4**

**ULP-3 is preferentially expressed in germ cells**. Thermo-sensitive mutants glp-4(bn2ts) do not develop germ line tissue into adult gonads when exposed to restrictive temperature at early development stage, leading to animals essentially devoid of germ cell as depicted in the upper panel. Wild type and glp-4(bn2ts) synchronised worms were both switched to restrictive temperature at the L1 larvae stage (25°C). After reaching young adult stage, worm protein were extracted and ULP-3 protein level analysed by western blotting (upper panel). RNA from whole animals or from dissected gonads from wild type animals was used for qPCR to monitor *ulp-3* expression. Values were normalised to tubulin expression and presented as fold change relative to whole animal *ulp-3* expression (lower right panel).

### Supplemental Figure 5

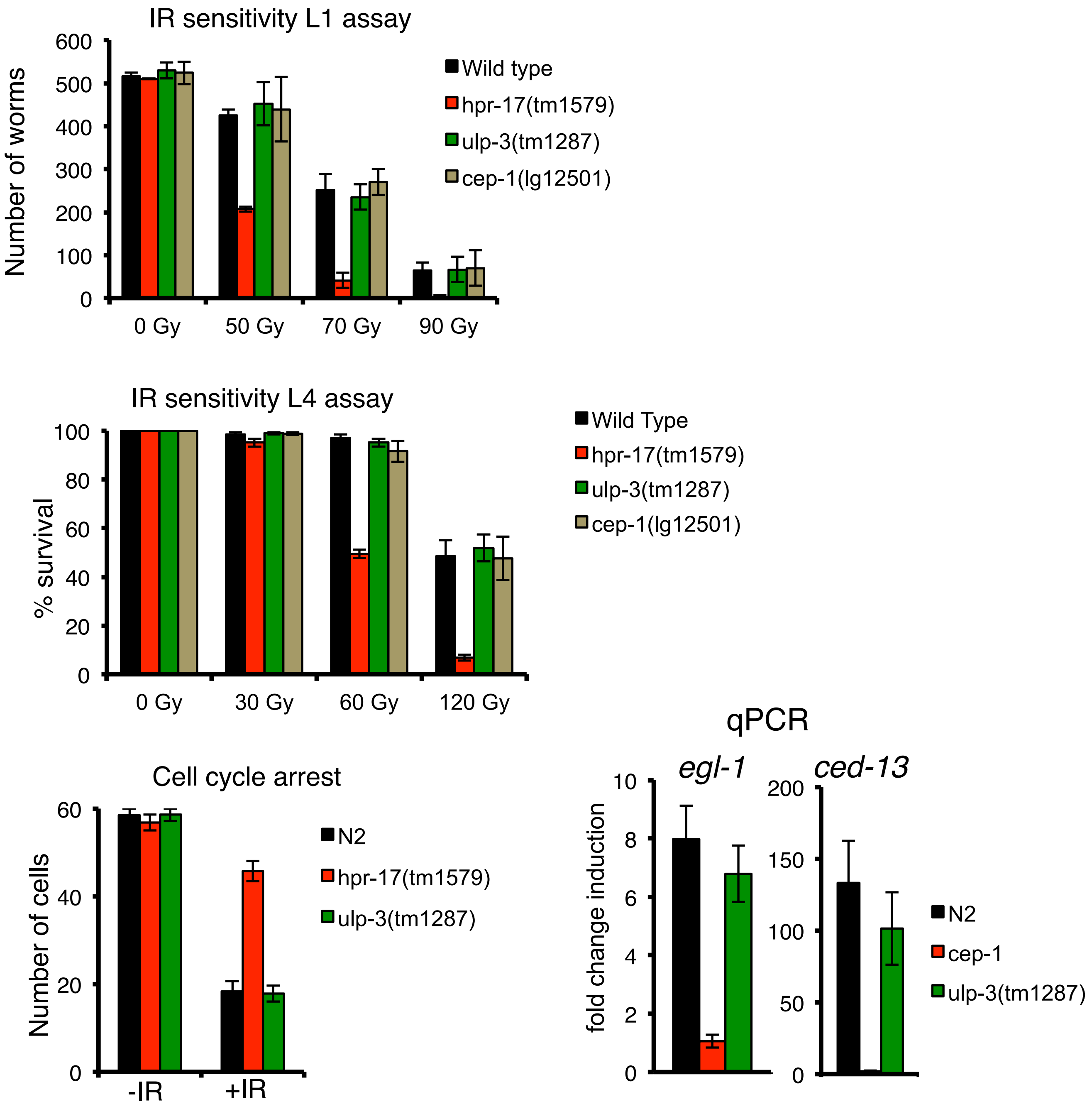


**Supplemental Figure 5**

***ulp-3* does not affect the DNA damage checkpoint pathway upstream of *cep-1***. The *ulp-3* deletion allele does not increase irradiation sensitivity following L1 (upper graph) or L4 (middle graph) larvae stage irradiation. Assays were performed as described (Bailly et al., 2010) using *hpr17* deletion mutant as a positive control (the RAD17 worm homologue). As shown in the lower graph *ulp-3* deletion does not impact on the mitotic cell cycle arrest triggered by the DNA damage checkpoint signalling pathway in *C. elegans* germ cells. Arrested cells were monitored and counted as previously described (Bailly et al., 2010). Expression levels of 2 p53 regulated genes (egl-1 and ced-13) after IR (90Gy) in different genetic backgrounds. γ-tubulin expression was used to normalise values. Data represent the mean of 3 independent experiments +/- SEM.

Supplemental Figure 6

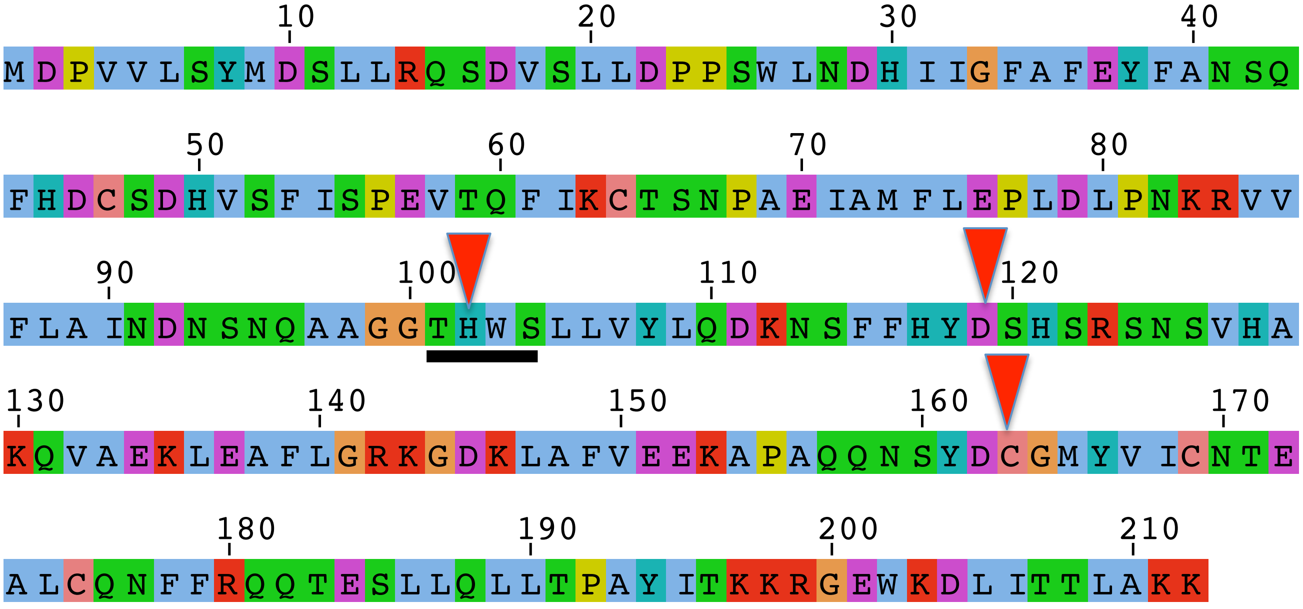


Parental 294-TGGAGGAACCCACTGGAGTTTATTGGTCTACCTCCAAGATA
H6 294-TGGAGGAACCC - - - - - - - - - - ATTGGTCTACCTCCAAGATA

**Supplemental Figure 6**
**NEDP1 deletion by CRISPR/Cas9.** Sequencing of NEDP1 PCR products using genomic DNA as template from parental or H6 U2OS cells confirmed the indicated deletion of 10 base pairs in the NEDP1 gene creating an out of frame sequence and a stop codon 90 bases downstream. Red arrows indicate the position of the catalytic triad in NEDP1 and the black line the target area of the guide sequence at amino acid level.

Supplemental Figure 7

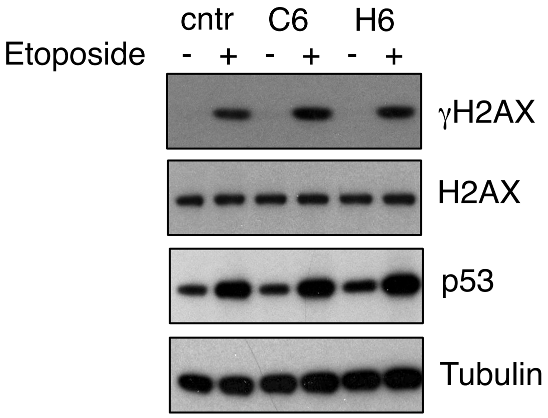


**Supplemental Figure 7**
**NEDP1 knockout does not affect γH2AX and p53 activation upon DSBs.** Parental or NEDP1 knockout U2OS cells (clones C6, H6) were treated with etoposide (50μM for 15hrs) and cell extracts were analysed with the indicated antibodies by Western blotting.

Supplemental Figure 8

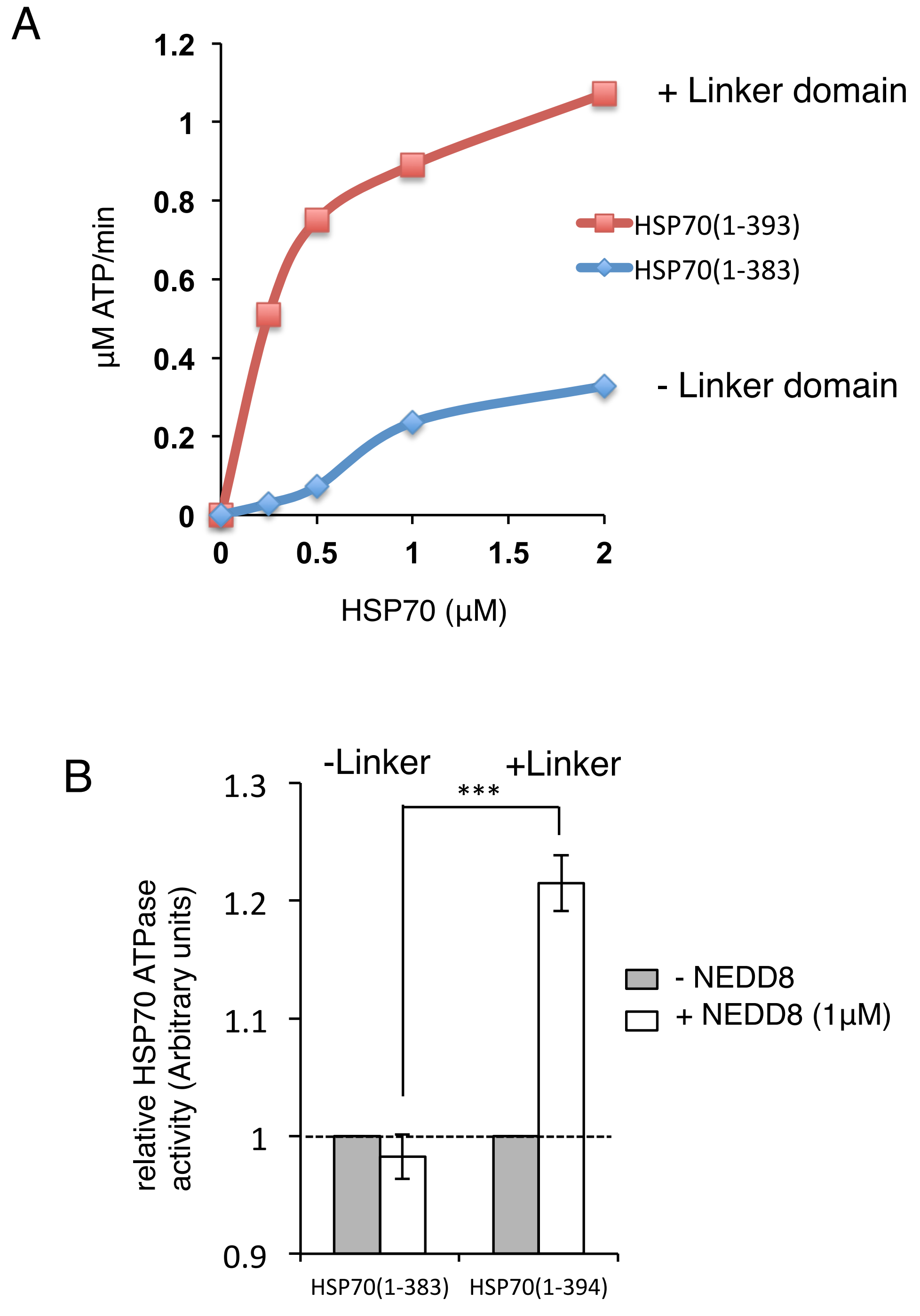


**Supplemental Figure 8**

**The Linker domain is critical for the stimulation of the HSP70 ATPase activity by NEDD8. A.** The ATP hydrolysis activity of the HSP70 ATPase domain in the presence or absence of the Linker domain (383-393) was measured. **B.** The effect of NEDD8 on the HSP70 ATP hydrolysis activity in the presence or absence of the linker domain in HSP70. Error bars represent SEM (*n*=5) with p value <0.001.

Supplemental Figure 9

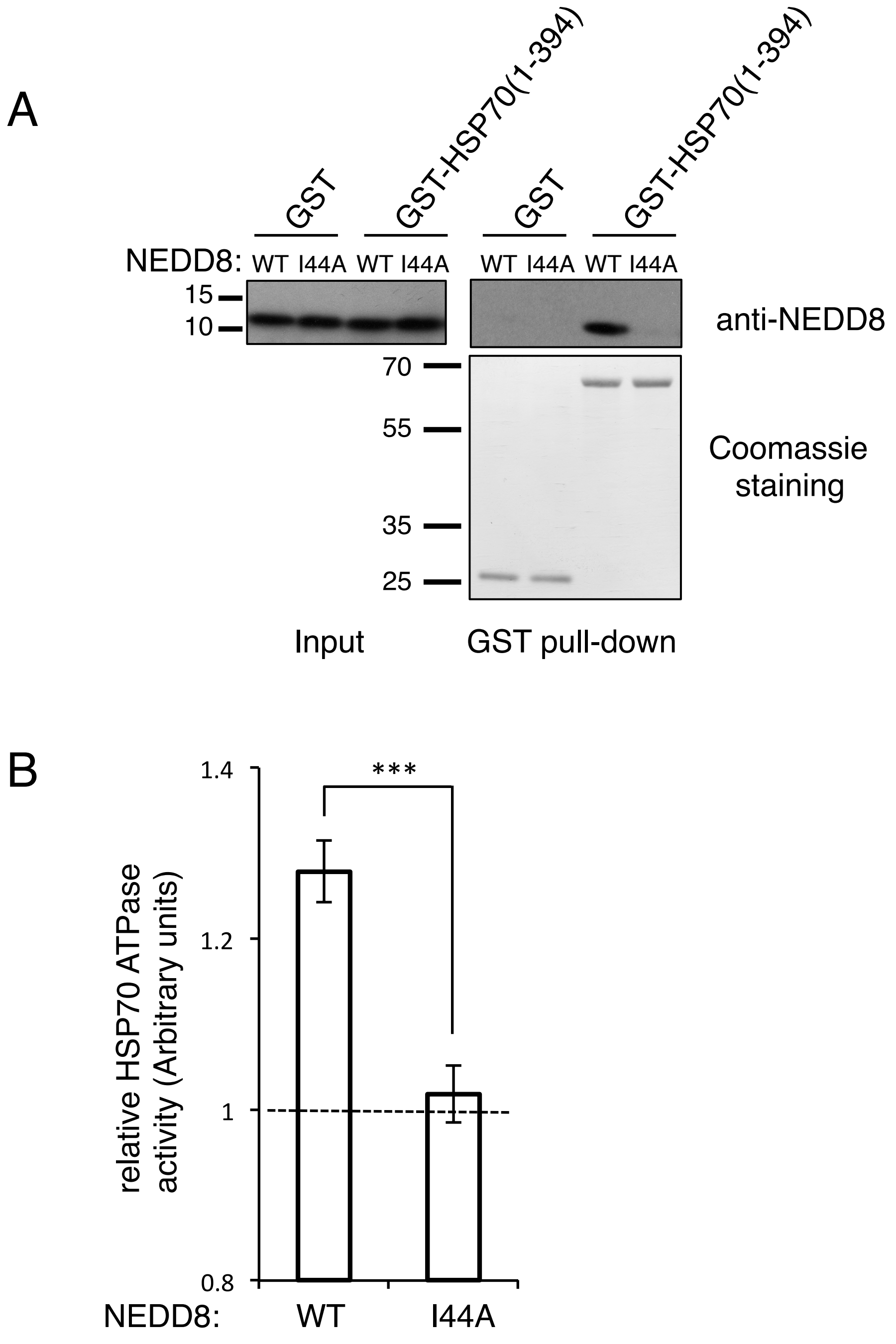

Supplemental Figure 9

C

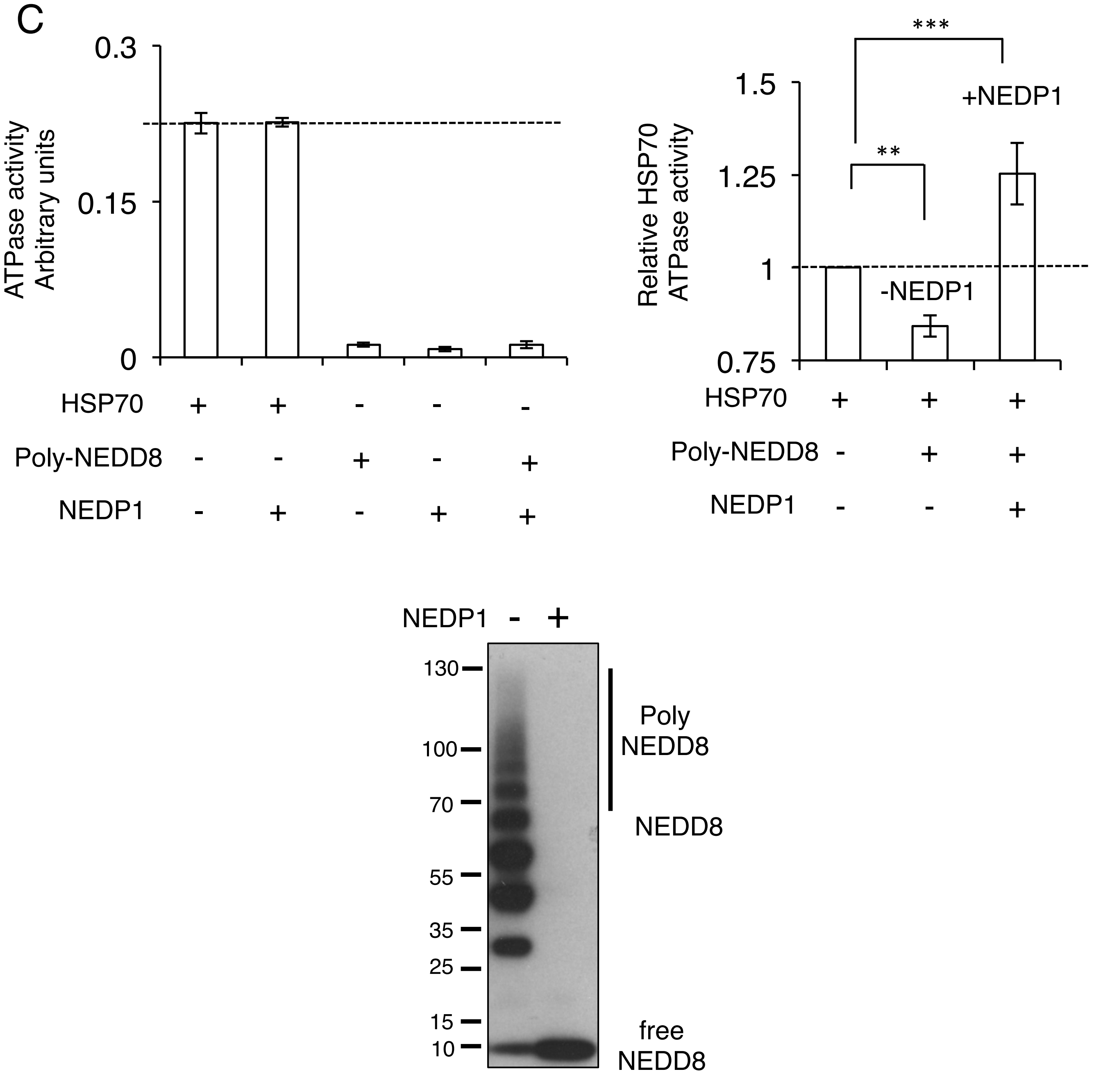


**Supplemental Figure 9**

**NEDD8 interacts with HSP70 ATPase domain through its hydrophobic patch residue I44. A.** The NEDD8 point mutant I44A fails to bind the HSP70 ATPase domain *in vitro* in a GST pull-down assay. **B.** The I44A mutation inhibits the NEDD8 stimulatory effect on HSP70 ATPase activity *in vitro*. Error bars represent SEM ($n$=5) with p value <0.001. **C.** Poly-NEDD8 conjugates synthesized *in vitro* using recombinant $His_6$-NEDD8 were isolated by $Ni^{2+}$ pull-down and subjected to NEDP1 digestion (bottom panel) and processed as in Fig.4F. ($n$=5 +/- SEM, p values **≤ 0.01, ***≤ 0.001).

**References**

Ellis, R.E., Jacobson, D.M., and Horvitz, H.R. (1991). Genes required for the engulfment of cell corpses during programmed cell death in Caenorhabditis elegans. Genetics *129*, 79–94.
